# Variation in microbial feature perception in the Rutaceae family with immune receptor conservation in citrus

**DOI:** 10.1101/2022.07.15.500235

**Authors:** Jessica Trinh, Tianrun Li, Jessica Y. Franco, Tania Y. Toruño, Danielle M. Stevens, Shree P. Thapa, Justin Wong, Rebeca Pineda, Emmanuel Ávila de Dios, Tracy L. Kahn, Danelle K. Seymour, Chandrika Ramadugu, Gitta L. Coaker

## Abstract

Although much is known about the responses of model plants to microbial features, we still lack an understanding of the extent of variation in immune perception across members of a plant family. In this work, we analyzed immune responses in *Citrus* and wild relatives, surveying 86 Rutaceae genotypes with differing leaf morphologies and disease resistances. We found that responses to microbial features vary both within and between members. Species in two subtribes, the Balsamocitrinae and Clauseninae, can recognize all tested microbial features (flg22, csp22, chitin), including one from *Candidatus* Liberibacter species (csp22_*C*Las_), the bacterium associated with citrus greening disease aka Huanglongbing. We investigated differences at the receptor level for flagellin perception (FLS2 receptor) and chitin perception (LYK5 receptor) in citrus genotypes. We were able to characterize two genetically linked *FLS2* homologs from ‘Frost Lisbon’ lemon (responsive) and ‘Washington navel’ orange (non-responsive). Surprisingly, *FLS2* homologs from responding and non-responding genotypes were expressed in citrus and functional when transferred to a heterologous system. ‘Washington navel’ orange weakly responds to chitin, but ‘Tango’ mandarin exhibits a robust response. LYK5 alleles were identical or nearly-identical between the two genotypes and able to complement the *Arabidopsis lyk4/lyk5-2* mutant with respect to chitin perception. Collectively, our data indicates that differences in chitin and flg22 perception in these citrus genotypes are not the result of sequence polymorphisms at the receptor level. These findings shed light onto the diversity of perception of microbial features and highlight genotypes capable of recognizing polymorphic pathogen features.

## Introduction

The perception of microbial features has typically been assessed by using a single or few plant genotypes to make conclusions about perception. Recognition of conserved features of pathogens, known as microbe-associated molecular patterns (MAMPs), activates the plant immune system. MAMPs can be proteinaceous or structural pathogen features and are perceived by plant immune receptors. While many studies are focused on the immune responses of one representative genotype, responses to MAMPs exhibit variation within and between related species. For example, different genotypes of *Arabidopsis thaliana* contain FLS2 homologs that vary in binding specificity to an epitope of bacterial flagellin, and low binding specificity correlated with high bacterial proliferation (Vetter *et al*. 2012). Epitopes from three bacterial MAMPs were differentially recognized across heirloom tomato genotypes, indicating the diversity in immune responses within a group of closely-related plants (Veluchamy *et al*. 2014). The extent of natural variation in immune responses to microbial features still remains largely unexplored.

The Rutaceae plant family contains ∼2100 species with worldwide distribution, including the agriculturally important genus *Citrus* (Kubitzki *et al*. 2011). Citrus is the most extensively produced fruit crop in the world with 124.246 million tons of fruit produced in 2016 (Zhong & Nicolosi 2020). In the United States, the 2019 - 2020 growing season yielded a value of production of about 3.4 billion dollars for citrus products (USDA 2020). Florida alone is the second largest producer of orange juice in the world behind Brazil, its citrus economy contributing billions of dollars to the state Gross Domestic Product (Hodges & Spreen 2012). While oranges constitute more than half of worldwide citrus production, other relevant citrus products include tangerines, limes, lemons, and grapefruits (Liu *et al*. 2012). These different varieties of cultivated citrus are members of the genus *Citrus* in the family of Rutaceae, which contains several non-cultivated relatives of *Citrus* (Wu *et al*. 2018). Several systems of classification exist; in this study, we followed the classification of Swingle and Reece (1967). Rutaceae contains six subfamilies (Appelhans *et al*. 2021) and the subfamily to which citrus belongs, Aurantoideae, contains two tribes: Citreae and Clauseneae (Morton 2009). The Citreae contains three subtribes, which are Citrinae, Triphasiinae, and Balasmocitrinae. Clauseneae contains three subtribes: Micromelinae, Clauseninae, and Merrillinae (Nagano *et al*. 2018). The Citrinae subtribe contains all cultivated citrus genotypes and is the most important group in the family Rutaceae (Swingle and Reece 1967).

Cultivated citrus is a perennial crop that is vegetatively propagated through grafting (Castle 2010, Caruso *et al*. 2020). While these asexual propagation methods maintain the desired combinations of traits in commercial cultivars, they prevent the exchange of genetic material (Uzun & Yesiloglu 2012, Wang *et al*. 2017). Because of this, crops that are primarily propagated asexually are susceptible to devastating impacts from newly introduced citrus diseases. Cultivated citrus varieties are susceptible to a variety of microbial pathogens including bacteria, filamentous pathogens, and viruses. Breeding efforts often focus on developing rootstocks with resistance to these pathogens to fend off disease in the clonally propagated scion. Examples of citrus diseases with a significant impact on citrus production include citrus Huanglongbing (HLB) (Bove *et al*. 2006, Wang 2019), citrus canker (Ference *et al*. 2018, Das 2003), citrus variegated chlorosis (Colleta-Filho *et al*. 2020), as well as fruit and root rots (Jaouad *et al*. 2020).

To protect themselves from pathogens, plants have evolved multiple defense mechanisms including MAMP perception. MAMPs can be proteinaceous, such as the flagellin or cold shock protein of bacteria (Wang *et al*. 2016) or not, such as bacterial lipopolysaccharide or fungal chitin (Newman *et al*. 2013). To detect MAMPs, plants possess pattern-recognition receptors (PRRs) on the surface of their cells. PRRs include receptor-like kinases (RLKs) and receptor-like proteins (RLPs). RLKs consist of an extracellular domain, a transmembrane domain, and an intracellular kinase domain, whereas RLPs lack the intracellular kinase domain. Examples of PRR extracellular domains include leucine-rich repeats (LRRs), lysine motifs (LysM), and lectin domains, among others (Ngou *et al*. 2022). Binding of the MAMP to the PRR often results in heterodimer formation with a coreceptor to activate downstream signaling responses and host defenses ultimately leading to MAMP-triggered immunity (MTI). Hallmarks of MTI activation include apoplastic reactive oxygen species (ROS) production, intracellular mitogen-activated protein kinase (MAPK) activation, calcium influx, and global transcriptional reprogramming (Saijo *et al*. 2017, Cuoto & Zipfel 2016, Bigeard *et al*. 2015, Jeworutzki *et al*. 2010).

Extensive research has been performed in the last few decades to reveal PRRs that perceive various MAMPs in model and crop plants (Ngou *et al*. 2022). Some well-characterized receptors include: the *Arabidopsis* FLS2 receptor for a 22-amino acid epitope of bacterial flagellin (Gómez-Gómez & Boller 2000), the *Arabidopsis* LysM domain receptor (LYK4/5) for chitin (Miya *et al*. 2007, Cao *et al*. 2014), and the tomato (*Solanum lycopersicum*) CORE receptor for a 22-amino acid epitope of bacterial cold shock protein (Wang *et al*. 2016). However, more work needs to be done to discover novel immune receptors in tree crops and other non-model plants. Genome mining for citrus LRR-RLKs suggest that there are receptors capable of mediating host-pathogen interactions. Although RLKs have been predicted in the *C. ×aurantium* L. and *C. clementina* genomes, no immune receptors have been functionally validated in citrus (Magalhães *et al* 2016, Dalio *et al*. 2017). Previous studies have identified citrus FLS2 homologs, one of which is induced in response to bacterial flagellin (Shi *et al*. 2016). In addition, one Liberibacter-specific MAMP for the bacterial protein pksG was recognized in three out of 10 citrus genotypes (Chen *et al*. 2020).

Here, we have examined the responses of several genotypes encompassing both cultivated citrus and wild relatives to different microbial features in order to better understand the landscape of immune perception within the Rutaceae family. FLS2 orthologs are present in both monocots and dicots. We identified nearly identical *FLS2* homologs from responding and non-responding citrus genotypes. Surprisingly, *FLS2* homologs from responding and non-responding genotypes were functional when transferred to a heterologous system, indicating that impaired flagellin perception is not due to differences at the receptor level. Most cultivated citrus and wild relatives can perceive chitin and we were able to isolate a citrus homolog of the *Arabidopsis* chitin receptor *LYK5* and demonstrate its functionality in a non-host species. We also identified citrus relatives that can perceive a conserved feature of *Candidatus* Liberibacter asiaticus (*C*Las), the bacterium associated with citrus Huanglongbing. These results highlight the importance of studying immunity in wild relatives, especially to identify potential genotypes with immune mechanisms that can be transferred to disease-susceptible cultivars.

## Results

### Members of the Rutaceae family exhibit diversity in the perception of and magnitude of response to microbial features

To investigate the immune response capabilities of members of the Rutaceae family, we screened 86 genotypes for the perception of three common microbial features: chitin, flg22, and csp22. These genotypes were samples from the Givaudan Citrus Variety Collection (GCVC) at the University of California, Riverside. The GCVC is one of the most comprehensive collections of citrus diversity, including over 1,000 accessions that span the genus *Citrus* and related genera. This study includes representatives spanning known subtribes in Rutaceae, including both cultivated citrus and wild relatives. In total, we screened individuals from two subfamilies (Aurantoideae and Zanthoxyloideae) and representative taxa from all six subtribes of Aurantoideae, comprising over 30 different genera. The majority of selected genotypes fall within the Citrinae subtribe (56 genotypes), which includes the cultivated citrus types. Genotypes are referred to by their common name, if available, with the corresponding scientific name and accession number in Table S1. To measure the ROS output of multiple genotypes, we have optimized a plate-based assay for high-throughput screening of leaves from both seedlings and mature trees. The genotypes we screened exhibit a variety of different leaf, branch, and fruit morphologies (Table S1, Fig. 1B). A luminol analog, L-012, chemically reacts with horseradish peroxidase and ROS to produce light, which is measured by the plate reader as relative light units (RLUs). RLUs from a ROS burst can be plotted as a curve over time, area under the curve, or in this case, the peak of ROS production (max RLU). The results from the ROS-based assay are presented as an average of max RLUs across multiple independent experiments in Fig. 1A.

**Figure 1.**
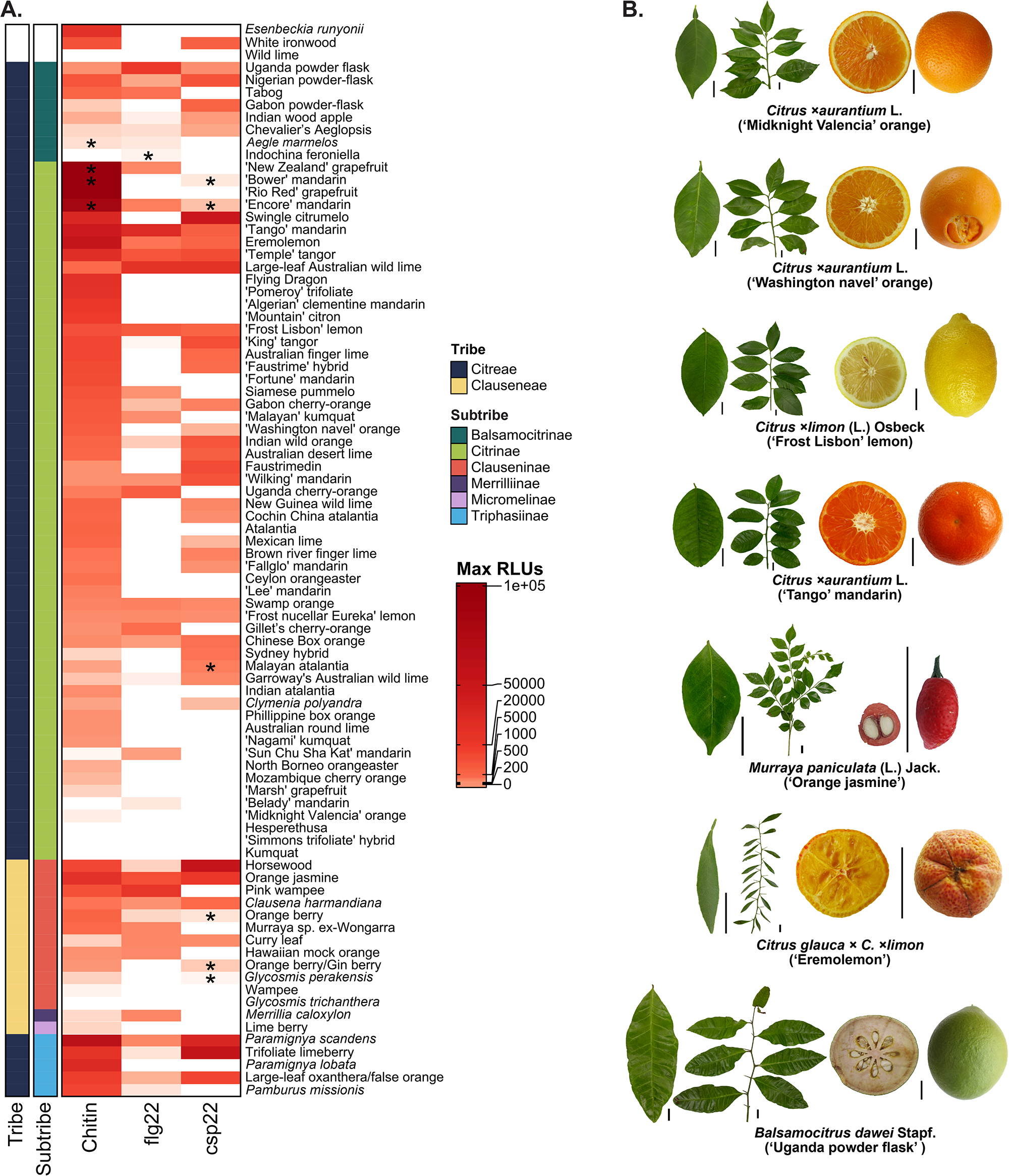
Genotypes within the Rutaceae family, including citrus, exhibit diverse responses to common MAMPs and possess differing leaf morphologies. A. Heat map compiling average max relative light units (RLUs) from reactive oxygen species (ROS) assays in genotypes within the Rutaceae family, organized by MAMP and phylogenetic relationship. Max RLUs are averages of at least three independent experiments and are represented as a heatmap, where *max RLU = (max RLU MAMP – max RLU water)*. <90 RLU is the threshold for no response. Asterisks indicate genotypes that exhibit a variable response, where one or two independent experiments shared a response. The MAMPs used are canonical features in the following concentrations: chitin (10 μM), flg22 (100 nM), csp22 (100 nM). B. Leaf, branch, and fruit morphologies of selected genotypes grown under greenhouse and field conditions. Scale bars = 2 cm. Genotypes are referred to by common name unless otherwise unavailable.

The landscape of immune perception varied across genotypes, with the strength of the response to each elicitor segregating across members from most surveyed tribes (Fig. 1A, Fig. S1). There were differences in the proportion of genotypes responding to each elicitor. For example, nearly all genotypes screened are capable inducing ROS in response to chitin (75 out of 86 genotypes), but less than half of the screened genotypes are capable of inducing ROS in response to flg22 (40 out of 86 genotypes). More than half of the screened genotypes (45 out of 86 genotypes) are capable of inducing ROS in response to csp22. The luminol assay used for ROS production can also be impacted by secondary compounds, including phenolics, antioxidants or reducing agents (Plieth 2018). The majority of genotypes were able to perceive chitin (76 out of 86), indicating secondary compounds in leaves did not grossly affect ROS production in these genotypes (Fig. 1, Fig. S1). Because of widespread chitin response across cultivated citrus and wild relatives, it is likely that these genotypes share a conserved chitin receptor, or multiple receptors capable of perceiving chitin of different lengths. There is substantial segregating variation in MAMP response across tribes. For example, within the Citrinae tribe, ‘Tango’ mandarin (*Citrus ×aurantium* L.) can respond to all three MAMPs, but kumquat (*Fortunella hindsii* (Champ. ex Benth.) Swingle) cannot respond to any of the three. Twenty-four genotypes are capable of responding to all three MAMPs in addition to ‘Tango’ mandarin. Closely related genotypes, such as ‘Tango’ mandarin and ‘Lee’ mandarin, also have varying responses to MAMPs: ‘Tango’ mandarin responds to all three MAMPs, but ‘Lee’ mandarin can only respond to chitin (Fig. 1A, Fig. S1). This variation may be the result of differences at the receptor level or in downstream signaling components.

Rutaceae genotypes vary not only in their ability to respond to MAMPs but also in the magnitude of ROS production (Figs. 1, 2, S1, S2). The magnitude of ROS production as a result of chitin, flg22, or csp22 induction occurs across tribes as well as within members of a tribe. Although the majority of the screened Rutaceae genotypes are capable of perceiving chitin, some genotypes produce an average max RLU of less than 1,000, while others produce an average max RLU well over 10,000 in response to chitin. To categorize the strength in responses, 25th and 75th quartiles were computed for each MAMP and cutoffs were used. Figure 2A shows examples of “weak” (25th percentile or below), “medium” (between the 25th and 75th percentiles), and “strong” (75th percentile or greater) responders. Across tribes, we see that members of the Triphasiinae tribe, such as the trifoliate limeberry, are strong ROS responders to chitin, whereas some members of the Balsamocitrinae subtribe, such as the Chevalier’s Aeglopsis, are either weak or medium ROS responders (Fig. 2B). Within the Citrinae tribe, ‘Tango’ mandarin and Uganda cherry-orange are strong responders to flg22, but ‘King’ tangor is a weak responder to flg22 (Fig. 2C). Trifoliate limeberry is also a strong responder to csp22 (Fig. 2D). The data showcase strong and weak ROS responders to MAMPs that are spread out within and between taxonomic groups.

**Figure 2.**
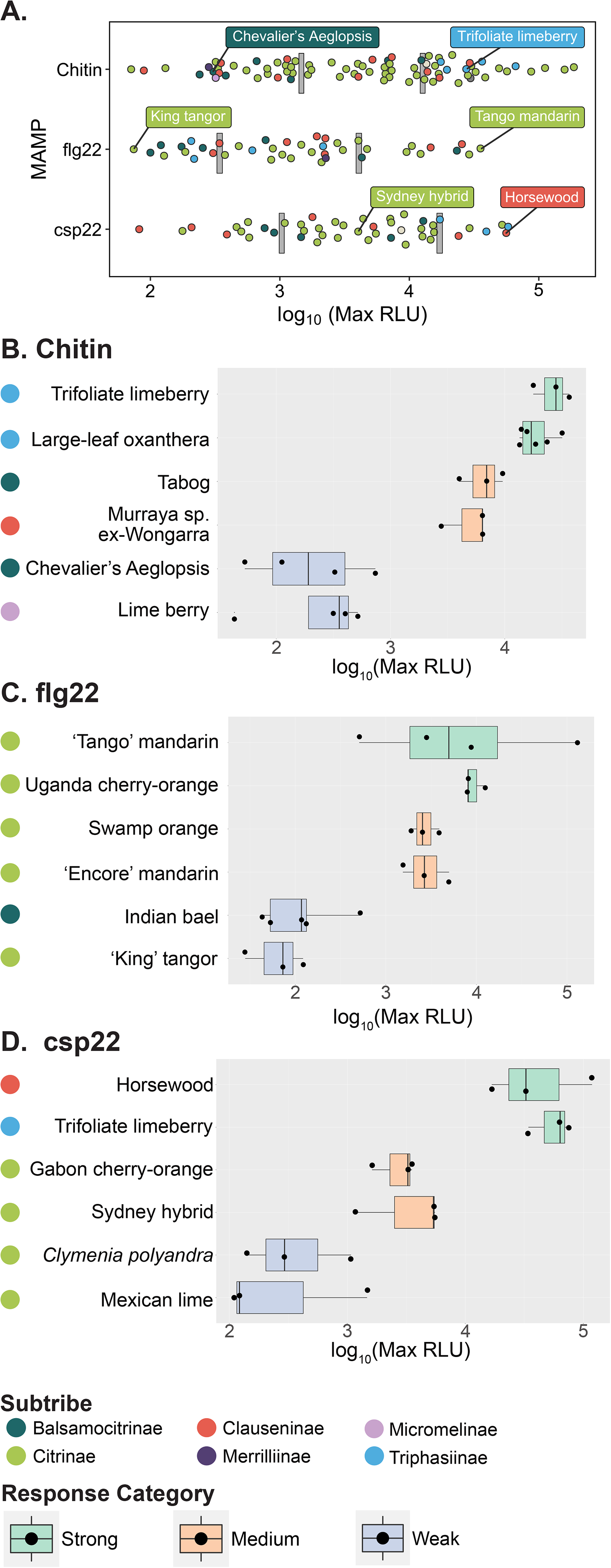
Rutaceae genotypes vary in the magnitude of their responses to perception of chitin, flagellin and cold shock protein immunogenic epitopes. A. Distribution of all average max RLU values, with gray lines indicating 25th and 75th percentile markings. Box plots below are organized by the magnitude of their responses to chitin (B), flg22 (C), and csp22 (D). The MAMPs used are canonical features in the following concentrations: chitin (10 μM), flg22 (100 nM), csp22 (100 nM). Max RLUs are averages of at least three independent experiments, where *max RLU = (max RLU MAMP – max RLU water)*. Data points on box plots represent the average max RLU for an individual experiment, with n = 8 leaf disks per experiment. Criteria for the response categories: “strong” responders are in the top 25th percentile, “medium” responders are between the 25th and 75th percentiles, and “weak” responders are in the bottom 25th percentile. The bar within the box plot depicts the median of the data, where the box boundaries represent the interquartile range (between the 25th and 75th percentiles) of the data. Box whiskers represent the minimum or maximum values of the data within 1.5× of the interquartile range.

In addition to the production of ROS, other common immune responses include MAPK activation, defense gene expression, and callose deposition. One of the challenges of studying Rutaceae is that many of the genotypes in this study do not have their genomes sequenced, making primer design for defense gene expression experiments difficult. Additionally, the thick, waxy leaves of citrus plants make it challenging to visualize callose deposition via microscopy. MAPKs are highly conserved across eukaryotes (Meng and Zhang 2013), making them viable immune markers to study MAMP responses in a variety of genotypes. MAPKs are phosphorylated upon MAMP perception, which can be detected via Western blot. MAPK phosphorylation can be weakly induced in response to water or buffer treatment but is strongly phosphorylated in response to immune activation (Asai *et al*. 2002, Zhang *et al*. 2013).

In order to investigate activation of other immune responses, we analyzed a subset of Rutaceae genotypes from three different subtribes (Balsamocitrinae, Clauseninae, and Citrinae) for PTI-induced MAPK activation in response to flg22 and chitin treatment. ‘Midknight Valencia’ orange is only able to respond to chitin based on ROS results and only exhibits MAPK phosphorylation upon chitin treatment (Fig. 3B). ‘Frost Lisbon’ lemon responds to chitin and flg22 based on ROS results and induces sustained MAPK phosphorylation in response to chitin and flg22 treatment. For Orange jasmine and Uganda powder flask, two non-Citrinae genotypes, we can observe ROS production in response to chitin and flg22. While chitin-induced MAPK phosphorylation was robust in both genotypes, flg22 perception was only observed in two out of four MAPK trials for the Uganda powder flask and Orange jasmine exhibited weak but reproducible MAPK phosphorylation after flg22 treatment. Taken together, these data indicate that Rutaceae genotypes can respond to MAMPs by inducing ROS and MAPK activation.

**Figure 3.**
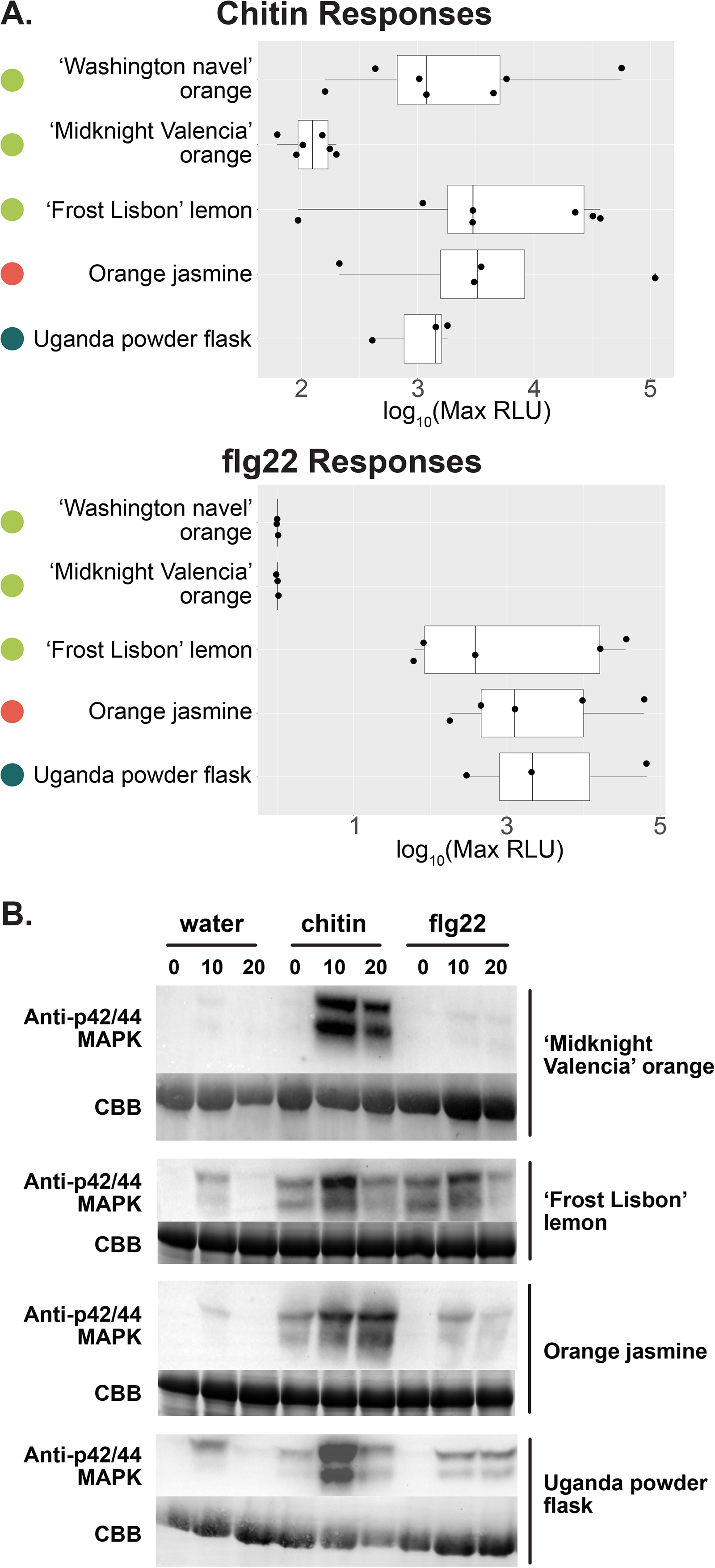
ROS and MAPK induction in response to MAMP treatment in cultivated citrus and wild relatives. A. Box plots showing the average max RLUs of selected Rutaceae genotypes in response to chitin and flg22. The MAMPs used are canonical features in the following concentrations: chitin (10 μM) and flg22 (100 nM). Max RLUs are averages of at least three independent experiments, where *max RLU = (max RLU MAMP – max RLU water)*. Data points on box plots represent the average max RLU for an individual experiment, with n = 8 leaf disks per experiment. The bar within the box plot depicts the median of the data, where the box boundaries represent the interquartile range (between the 25th and 75th percentiles) of the data. Box whiskers represent the minimum or maximum values of the data within 1.5× of the interquartile range. B. MAPK induction visualized at 0, 10, and 20 min post-induction with water or MAMP. MAMPs are applied to leaf punches at the following concentrations for MAPK assays: chitin (10 μM), flg22 (100 nM), with water as a negative control. Western blots are performed with an anti-p42/44 MAPK antibody to visualize the MAPK bands and Coomassie Brilliant Blue (CBB) to verify equal loading of protein samples. All experiments were performed at least 3X; flg22 perception was not observed in 2 out of 4 trials for the Uganda powder flask.

### Functional analyses of FLS2 in flagellin-responsive and nonresponsive citrus genotypes

*FLS2* orthologs have been identified and functionally validated from diverse plant families including Brassicaceae (Gómez-Gómez et al. 2000), Solanaceae (Robatzek et al. 2007), Vitaceae (Trdá et al. 2014) and Poaceae (Takai et al. 2008). Of the 86 genotypes we surveyed, 41 were able to perceive flg22 (Fig. 1A, S1). Sweet orange genotypes, including ‘Midknight Valencia’ orange and ‘Washington navel’ orange, were not able to elicit a ROS response to flg22, unlike the cultivated lemon genotypes ‘Frost Lisbon’ lemon and “Frost nucellar Eureka’ lemon (Fig. 1A, Fig. 4A). ‘Washington navel’ orange and ‘Frost Lisbon’ lemon are two widely grown citrus genotypes. Therefore, we investigated their response to flg22 in more detail. MAPK assays after treatment with flg22 verified that ‘Washington navel’ orange could not respond, while ‘Frost Lisbon’ lemon was able to induce robust MAPK phosphorylation (Fig. 4B). Similarly, flg22 treatment induced expression of the defense marker gene *WRKY22* in ‘Frost Lisbon’ lemon but not ‘Washington navel’ orange (Fig. 4C). Previously, two *FLS2* homologs (*FLS2-1* and *FLS2-2*) from ‘Duncan’ grapefruit and ‘Sun Chu Sha Kat’ mandarin were identified and demonstrated to be genetically linked (Shi et al. 2016). Similarly, when we analyzed the genomes of ‘Washington navel’ orange and ‘Frost Lisbon’ lemon, we identified *FLS2-1* and *FLS2-2* in syntenic chromosomal regions on haplotype 2 (Fig. 4D, Fig S3). Interestingly, *FLS2-2* is absent in ‘Frost Lisbon’ lemon haplotype 1 and truncated in ‘Washington navel’ orange haplotype 1 (Fig. 4D). Due to high conservation between *FLS2-1* alleles from each haplotype, we were unable to distinguish their transcripts by qPCR. Both genotypes exhibited low, but detectable baseline expression of each homolog using qPCR (Fig. S4). *FLS2-1* and *FLS2-2* had higher baseline transcript expression in ‘Washington navel’ orange, indicating differential responsiveness is not due to resting-state expression (Fig. S4). In both citrus genotypes, *FLS2-2* expression was induced after treatment with flg22, with ‘Frost Lisbon’ lemon exhibiting stronger induction (Fig. 4E). The immune responses (ROS, MAPK) we measured occur within 10 minutes post-MAMP treatment and both citrus genotypes express *FLS2-1* and -*2* in the absence of flg22 perception. Therefore, it is unlikely that differences in early immune outputs would be regulated by *de novo* transcription of *FLS2* PRRs.

**Figure 4.**
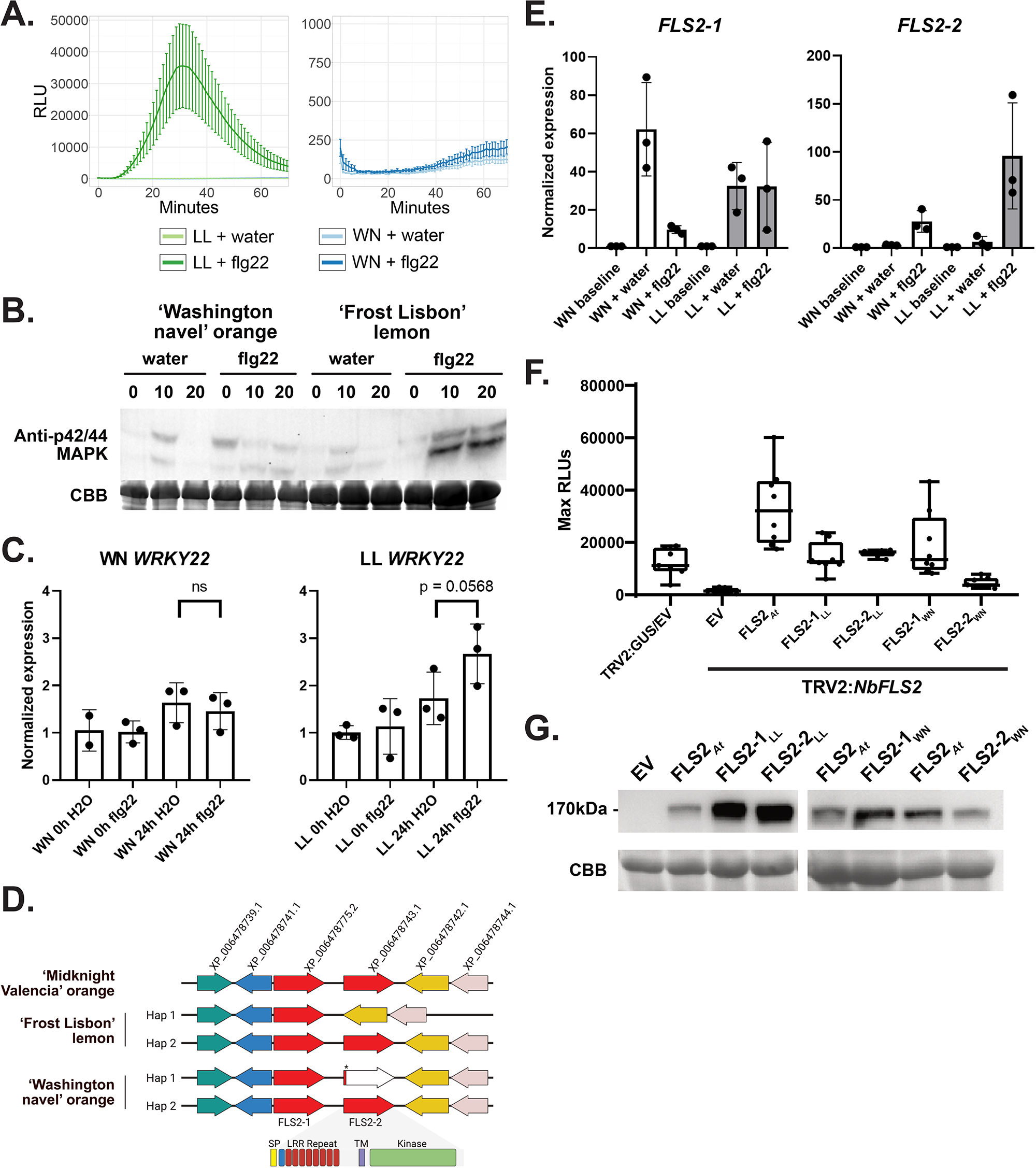
‘Washington navel’ orange (WN) and ‘Frost Lisbon’ lemon (LL) have nearly identical FLS2 homologs but differ in their response to bacterial flagellin. A. ROS curve for LL (left) and WN (right) when induced with either water or 100 nM flg22. Note the different scale on the y-axes. B. MAPK induction in response to either water or flg22 in WN vs. LL using anti-p42/44 MAPK immunoblotting. Experiments were repeated 3 times. CBB = Coomassie Brilliant Blue. C. Normalized expression of *WRKY22* after induction with either water or 10 μM flg22. Significance was determined via one-way ANOVA with a Sidak’s multiple comparisons test to determine significance between water and flg22 treatments at 24h. Bullet points represent technical replicates from a single tree. Experiments were repeated 4X with similar results. D. Genome organization of *FLS2* homologs in citrus, with the two chromosomally linked homologs highlighted in red. Haplotype data are shown for ‘Frost Lisbon’ lemon and ‘Washington navel’ orange. E. Expression of *FLS2-1* and *FLS2-2* transcripts measured via qPCR at resting state and when induced with either flg22 or water, using citrus *GAPDH* as a reference gene. Error bars represent the standard deviation (n = 3 biological replicates). Note the different axes scales between *FLS2-1* and *FLS2-2*. F. Transcomplementation experiments for FLS2 function in *Nicotiana benthamiana*. Two to three weeks post-silencing of *N. benthamiana* with tobacco rattle virus (TRV) targeting *GUS* (negative control) or endogenous *FLS2* (TRV2:NbFLS2), *FLS2* homologs from different plants were expressed using *Agrobacterium*-mediated transient expression. Forty-eight h post-*Agrobacterium* infiltration, leaf disks were subjected to ROS burst assays after treatment with 100 nM flg22, error bars = standard deviation, n > 7. G. Western blot demonstrating expression of all FLS2 proteins 48h post-*Agrobacterium* infiltration. Top: anti-HA-HRP blot, bottom: Coomassie brilliant blue (CBB) staining. EV = empty vector, _*At*_ = *Arabidopsis thaliana*.

To test if compromised flagellin perception in ‘Washington navel’ orange is due to sequence polymorphisms in *FLS2*, we investigated the ability of each homolog to perceive flg22 using transcomplementation experiments in *Nicotiana benthamiana.* We used Virus-Induced Gene Silencing (VIGS) to silence endogenous *FLS2* in *N. benthamiana,* followed by *Agrobacterium*-mediated transient expression of *Arabidopsis FLS2* as well as citrus *FLS2-1* and *FLS2-2*. Forty-eight hours after transient expression, we assayed silenced plants for their ability to induce a ROS burst in response to flg22 treatment. As expected, *N. benthaminana FLS2* silenced lines were unable to elicit a flg22-induced ROS burst, but *GUS* silenced lines were able to perceive flg22 (Fig. 4F). Expression of *Arabidopsis* FLS2, ‘Frost Lisbon’ lemon FLS2-1 and FLS2-2 led to a ROS burst in response to flg22 and detectable using anti-HA western blotting (Fig. 4F-G). Expression of ‘Washington navel’ orange FLS2-1, and to a lesser extent, FLS2-2, was also able to elicit a ROS burst in response to flg22 (Fig. 4F). While ‘Washington navel’ orange FLS2-1 was robustly expressed by western blot, FLS2-2 exhibited lower level expression, which may explain its reduced ROS burst (Fig. 4G). These data are consistent with the near identical amino acid similarity between FLS2-1 (99.16-99.41%) and FLS2-2 (97.41%) (Fig. S3). Collectively, these results suggest that differences in flg22-mediated responses between both genotypes may not be regulated at the receptor level.

### Cultivated citrus genotypes contain functional chitin receptor homologs

In our experiments, chitin is widely perceived across members of the Rutaceae, including both cultivated citrus types and wild relatives (Fig. 1A, S1), indicating that chitin perception is likely derived from a conserved receptor. Therefore, we sought to further investigate the presence of *LYK5,* the major chitin receptor (Xue *et al*. 2019, Erwig *et al*. 2017, Cao *et al*. 2014), and *CERK1* across the plant kingdom. To identify and assess the conservation of *Arabidopsis thaliana* LYK5_*At*_ and CERK1_*At*_ across a variety of eudicots and monocots, we used an approach based on homology, phylogeny, and hidden Markov models. Mid-rooted maximum likelihood trees show broad conservation of LYK5 and CERK1, with 41% of genotypes possessing multiple LYK5_*At*_ homologs and 51% possessing multiple CERK1_*At*_ homologs (Fig. 5A). There are two predominant LYK5 clades, a monophyletic monocot clade and a polyphyletic dicot clade including members from the Brassicaceae, Fabaceae, Malvaceae, and Rutaceae families. CERK1 displayed two major clades split by homologs from eudicots and monocots. Despite the diversification that can be found within the ectodomain of LYK5 and CERK1 homologs when compared against *Arabidopsis*, residues in LYK5_*At*_ which are known to directly bind to chitin are conserved (Fig. 5B-C) (Cao *et al*. 2014).

**Figure 5.**
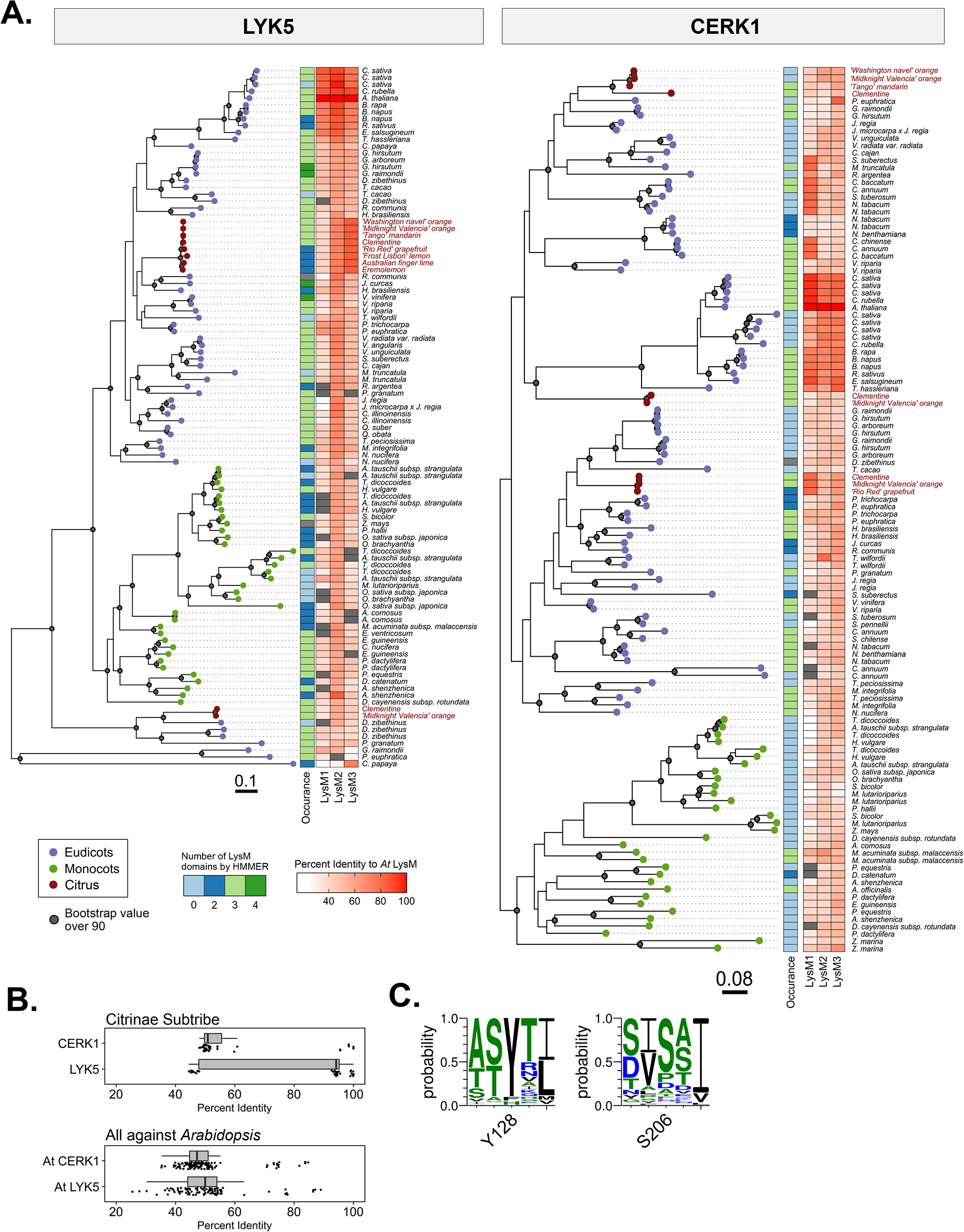
Phylogeny of LYK5 and CERK1 receptor homologs. A. Maximum likelihood phylogenetic tree of 102 LYK5 and LYK5-like homologs (top) and 120 CERK1 and CERK1-like homologs (bottom) from 66 plant species. In both trees, eudicots are labeled in purple, monocots are labeled in green, and sequences from citrus varieties are labeled in red. 1000 ultrafast bootstrap replicates were calculated and values over 90 were plotted as a gray dot. To determine the number of LysM domains, hmmer (LysM domain, query ID: PF01476.19) and blastp was used. Similarity to the *Arabidopsis thaliana* LysM domains by blastp from LYK5 and CERK1 were calculated and plotted. Scale bar indicates tree distance. B. All-by-all blastp of LysM receptor ectodomains. Top: blastp comparison of LYK5 and CERK1 homologs from the Citrinae tribe. Bottom: blastp comparison of all LYK5 and CERK1 plant homologs using *Arabidopsis* as a query. C. Weblogos across 102 plant LYK5 homologs corresponding to critical residues for chitin binding in *A. thaliana* LYK5_*At*_ Y128 and S206.

Additional citrus LYK5 members for Australian finger lime, ‘Rio Red’ grapefruit, ‘Frost Lisbon’ Lemon and Eremolemon were PCR amplified, sequenced, and plotted on the phylogeny. LYK5 homologs within the Citrinae tribe exhibit low copy number and diversification, predominantly clustering in a single subclade (Fig. 5A). We attempted to PCR amplify additional *CERK1* homologs in Citrinae but were only able to amplify from ‘Rio Red’ grapefruit. Within eudicots, citrus CERK1 homologs are polyphyletic and both clementine and ‘Midknight Valencia’ orange carry multiple homologs (Fig. 5A). Ectodomains of citrus receptor homologs were compared, revealing a bimodal distribution for LYK5 and CERK1 and high amino acid similarity within citrus subclades (Fig. 5B).

Although most cultivated citrus genotypes can respond to chitin, there is still a wide range for the magnitude of the ROS response (Fig. 1A). ‘Washington navel’ orange (*Citrus ×aurantium* L.) is a sweet orange and many modern type III mandarins are often derived from hybrids of sweet oranges and other mandarin types (Wu *et al*. 2018). ‘Tango’ mandarin has a stronger response to chitin, with a five-fold stronger ROS burst (Fig. 6A). Chitin also activates MAPKs in both ‘Washington navel’ orange and ‘Tango’ mandarin, though the magnitude of the response varies in ‘Tango’ mandarin (Fig. 6B). Similarly, chitin treatment more robustly induced expression of the defense marker gene *WRKY22* in ‘Tango’ mandarin compared to ‘Washington navel’ orange (Fig. 6C). We identified *LYK5* alleles on each chromosome of ‘Washington navel’ orange and ’Tango’ mandarin (type III, Fig. S5). Both genotypes contain homologous *LYK5* genes in syntenic genetic regions that are highly similar to each other with conserved chitin binding residues (99.6% amino acid similarity, Fig. 6D, S5). Moreover, ‘Washington navel’ orange and ‘Tango’ mandarin possess one identical allele of *LYK5* (allele 1) and a second nearly identical allele with two amino acid polymorphisms (allele 2, Fig. S5). One polymorphic residue is between transmembrane and kinase domain, while the second is in the kinase domain but not in a known critical residue (Fig. S5). These *LYK5* homologs are also expressed similarly in both genotypes using qPCR (Fig. 6E).

**Figure 6.**
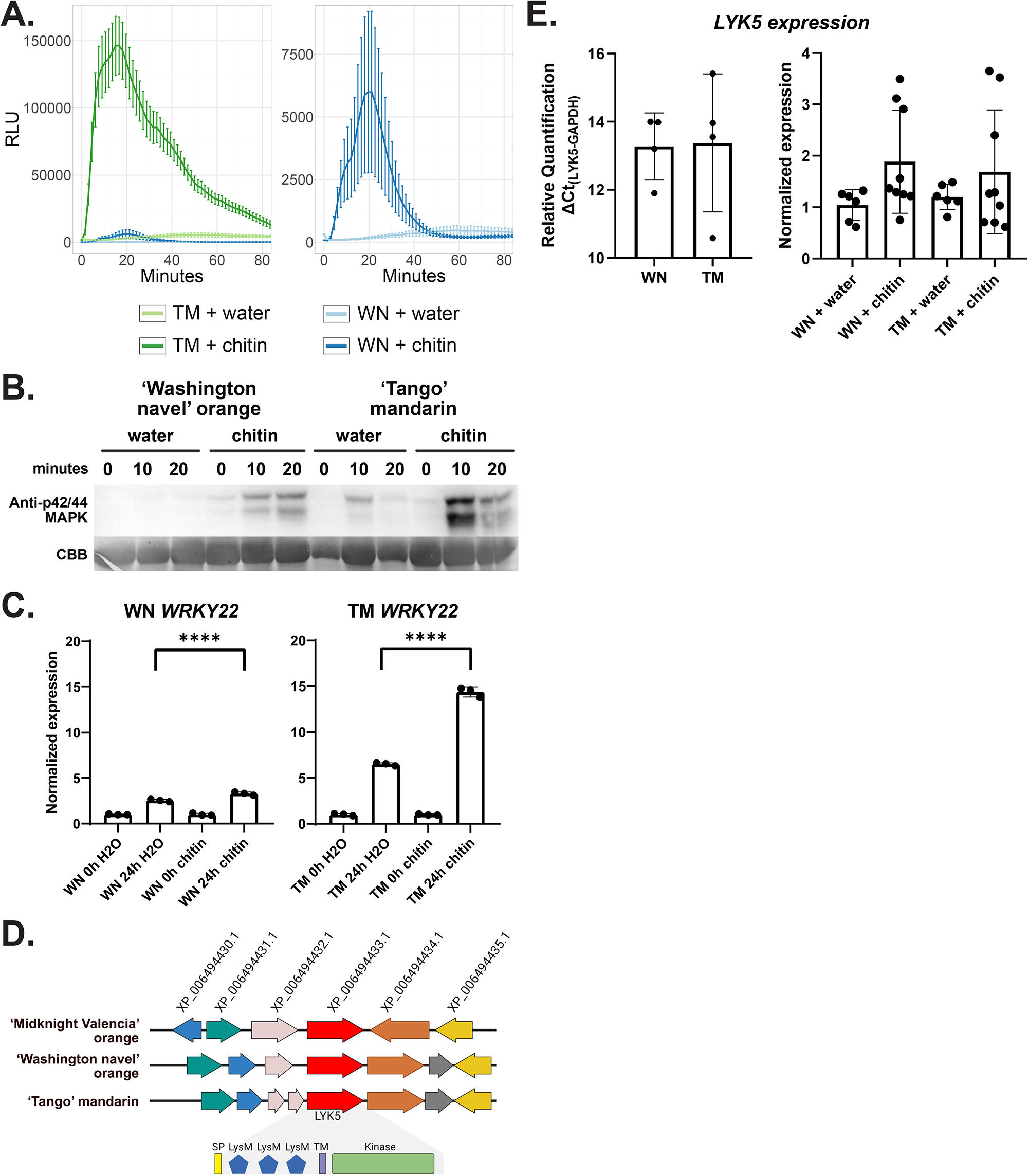
‘Washington navel’ orange (WN) and ‘Tango’ mandarin (TM) have nearly identical LYK5 homologs but differ in magnitude of their chitin response. A. ROS curve for ‘Washington navel’ orange and ‘Tango’ mandarin (left) and ‘Washington navel’ orange only (right) when induced with either water or 10 μM chitin. Note the different scale on the y-axes. B. MAPK induction in response to either water for chitin in ‘Washington navel’ orange vs. ‘Tango’ mandarin using anti-p42/44 MAPK immunoblotting. Experiments were repeated 4 times. CBB = Coomassie Brilliant Blue. C. Normalized expression of *WRKY22* after induction with either water or 10 μM chitin. Significance was determined via one-way ANOVA with a Sidak’s multiple comparisons test to determine significance between water and chitin treatments at the 24-hour mark. Bullet points represent technical replicates from a single tree. Experiments were repeated 4X; significant ‘Washington navel’ orange induction with chitin was only observed in 2 out of 4 trials. D. Genome organization of *LYK5* in ‘Midknight Valencia’ orange, ‘Washington navel’ orange, and ‘Tango’ mandarin, with the LYK5 domain in ‘Tango’ mandarin expanded to show functional domains. Arrows indicate the difference in amino acid sequence between ‘Tango’ mandarin and ‘Washington navel’ orange. E. Transcript expression of citrus *LYK5* measured via qPCR at resting state, using citrus *GAPDH* as a reference gene. Error bars represent the standard deviation (n = 4 biological replicates). E. Transcript expression of citrus *LYK5* transcript via qPCR at resting state (left, ΔCt) and when induced with water or chitin (right), using citrus *GAPDH* as a reference gene. Error bars represent the standard deviation (n = 4 biological replicates).

To validate the functionality of *LYK5* allele 1, we complemented *lyk4/lyk5-2 A. thaliana* with the ‘Tango’ mandarin LYK5 (identical to ‘Washington navel’ orange allele 1, referred to as LYK5_™_) or *Arabidopsis* LYK5_*At*_. Two independent *A. thaliana* transgenic lines expressing LYK5_™_ as well as LYK5_*At*_ regain the ability to produce ROS in response to chitin treatment (Fig. 7A). Immunoblot analyses against the HA epitope tag verified protein expression of all transgenes (Fig. 7B). We were able to further confirm the functionality of this receptor with MAPK assays. Both LYK5_™_ as well as LYK5_*At*_ complementation lines exhibited MAPK phosphorylation upon chitin treatment (Fig. 7C). Taken together, we have demonstrated that cultivated citrus can respond to chitin and possess *LYK5* and *CERK1* homologs. The LYK5 allele 1 can also function as a chitin receptor in *Arabidopsis*.

**Figure 7.**
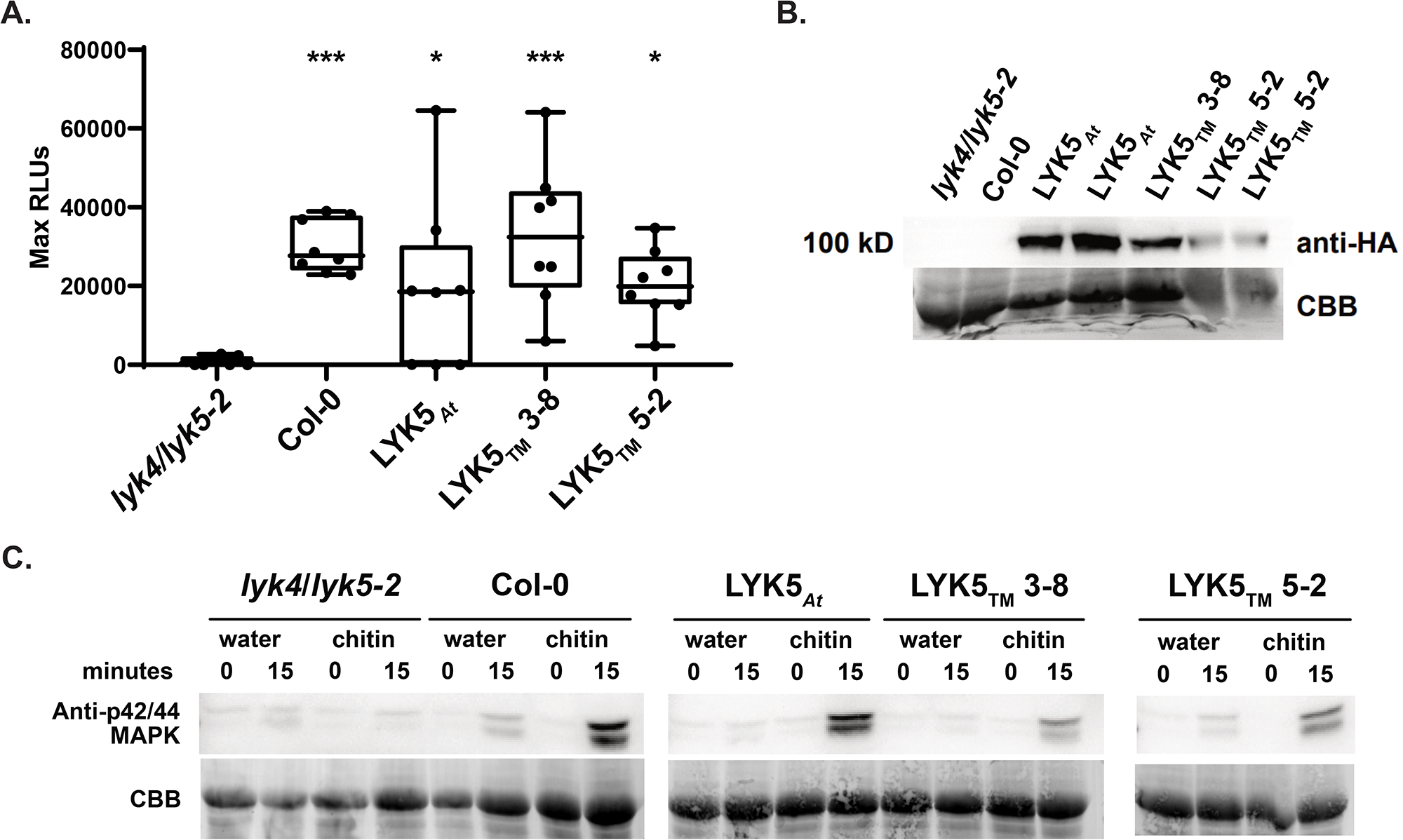
The ‘Tango’ mandarin LYK5_*At*_ homolog can complement an *Arabidopsis* chitin perception mutant. A. ROS production of *Arabidopsis lyk4/lyk5-2* knockouts complemented with the indicated LYK5 constructs after treatment with 10 μM chitin. We complemented *Arabidopsis* with *LYK5* allele 1, which is referred to as *LYK5_™_* and is identical between ‘Tango’ mandarin and ‘Washington navel’ orange. The bar within the box plot depicts the median of the data, where the box boundaries represent the interquartile range (between the 25th and 75th percentiles) of the data. Box whiskers represent the minimum or maximum values of the data within 1.5× of the interquartile range. Significance of results was determined via ordinary one-way ANOVA, with a post hoc Dunnett’s multiple comparison to the *lyk4/lyk5-2* knockout. Asterisks represent significance thresholds: *** p=0.0001 to 0.001, * p=0.01 to 0.05. B. Anti-HA-HRP immunoblots visualize the expression of LYK5-HA homologs in *Arabidopsis*; CBB = Coomassie Brilliant Blue (CBB). C. MAPK induction in response to either water for chitin in LYK5-complemented *Arabidopsis lyk4/lyk5-2*, using anti-p42/44 MAPK immunoblotting. CBB = Coomassie Brilliant Blue. All experiments were performed 3 times with similar results.

### Members from the Citrinae, Balsamocitrinae, and Clauseninae subtribes are capable of perceiving csp22 from an important citrus pathogen

In tomato, the CORE RLK perceives csp22, generating resistance to the bacterial pathogen *Pseudomonas syringae pv. tomato* DC3000 when expressed in *A. thaliana* (Wang *et al*. 2016). The closest *N. benthamiana* homolog of tomato CORE is able to induce ROS production in response to csp22 treatment after transient expression (Wang *et al*. 2016). In order to gain insight into candidate citrus csp22 receptors, we analyzed citrus genomes for the presence of CORE receptor homologs. However, the closest citrus homolog of either *Nicotiana* or *Solanum* CORE receptors had limited sequence similarity (Fig. S8). Expression of the receptor recognizing csp22 is developmentally-regulated and expressed in flowering *N. benthamiana* and tomato (Saur *et al*. 2016, Wang *et al*. 2016), which makes it possible to use *Agrobacterium*-mediated transient expression to investigate CORE in young *N. benthamiana*. We investigated the CORE homologs from ‘Frost nucellar Eureka’ lemon and ‘Washington navel’ orange, which are responsive to csp22. When these CORE homologs were heterologously expressed in 30 day-old *N. benthamiana*, they failed to confer csp22 responsiveness, in contrast to expression of *NbCORE*. All CORE proteins were expressed by immunoblot analysis (Fig. S8C). These data suggest that Rutaceae possess an independently derived csp22 receptor.

Proteinaceous MAMPs are often conserved across pathogens. However, due to strong selection pressure, some pathogens have evolved immunogenic epitopes that cannot be perceived, while still retaining the presence of the entire protein (Cheng *et al*. 2021). The csp22 epitope from *C*Las (csp22_*C*Las_) contains several polymorphisms when compared to the canonical csp22 sequence (Fig. 8A). Therefore, we investigated if there were members of the Rutaceae family that could perceive csp22_*C*Las_ (Fig. 8B). The vast majority of Rutaceae genotypes that can respond to canonical csp22 cannot perceive csp22_*C*Las_ using ROS production as an output. Notably, members from the Balsamocitrinae and Clauseninae subtribes can perceive both canonical csp22 and csp22_*C*Las_, such as Uganda powder-flask and *Clausena harmandiana*. There is one member of the Citrinae tribe, the ‘Algerian clementine’ mandarin, that perceives csp22_*C*Las_, but not canonical csp22. *Clausena harmandiana* is able to induce MAPK phosphorylation in response to csp22_*C*Las_ compared to the non-responding genotype ‘Midknight Valencia’ orange (Fig. 8D). Genotypes that respond to csp22_*C*Las_ also exhibited some level of reduced symptomology to HLB disease in field trials with mature trees (Fig. 8, S7, Ramadugu *et al*. 2016). However, not all genotypes with HLB tolerance can respond to csp22_*C*Las_ (Fig. S7). Data generated from our ROS screens in members of Rutaceae reveal members that can be used to identify novel receptors for transfer to *C*Las-susceptible citrus cultivars.

**Figure 8.**
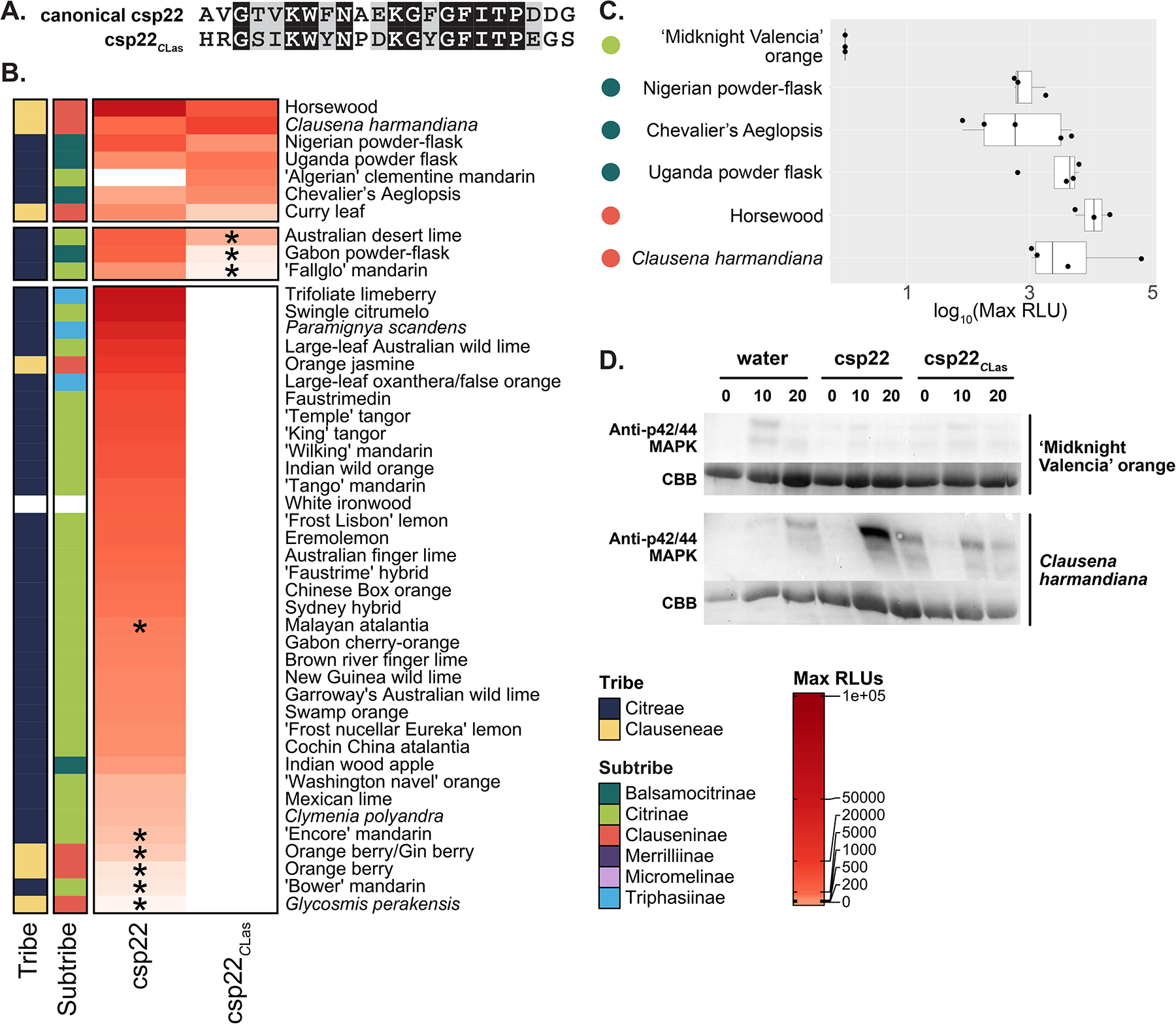
Three Rutaceae tribes can respond to a polymorphic csp22 from an important citrus pathogen. A. Alignment of the canonical csp22 sequence to the csp22 of *Candidatus* Liberibacter asiaticus (csp22_*C*Las_). B. Heat map compiling average max relative light units (RLUs) from ROS assays in genotypes within the Rutaceae family, organized by MAMP and phylogenetic relationship. Max RLUs are averages of at least three independent experiments, where *max RLU = (max RLU MAMP – max RLU water)*. <90 RLU is the threshold for no response. Asterisks indicate genotypes that exhibit a variable response, where one or two independent experiments revealed a response. The MAMPs used for treatments are canonical csp22 and csp22 from *Candidatus* Liberibacter asiaticus. C. Box plot of max RLUs for citrus relatives that can respond to 200 nM csp22_*C*Las_. Max RLUs are averages of at least three independent experiments. Data points on box plots represent the average max RLU for an individual experiment, with n = 8 leaf disks per experiment. The max RLU are plotted on a log10 scale. The bar within the box plot depicts the median of the data, where the box boundaries represent the interquartile range (between the 25th and 75th percentiles) of the data. Box whiskers represent the minimum or maximum values of the data within 1.5× of the interquartile range. D. MAPK induction by csp22 or csp22_*C*Las_ in either ‘Midknight Valencia’ orange or *Clausena harmandiana*, visualized by p42/44 MAPK antibody immunoblotting. CBB = Coomassie Brilliant Blue.

## Discussion

Here, we have investigated variation in MAMP perception within the Rutaceae family to determine the landscape of perception in citrus and citrus relatives. Variations in MAMP perception have been noted within genotypes of the same species, such as in tomato (Roberts *et al*. 2019) and in *Arabidopsis* (Vetter *et al* 2012). Even in close relatives, the perception of a potent immune elicitor such as flg22 varies widely (Veluchamy *et al*. 2014). Much more diversity remains to be discovered by analyzing multiple genotypes. Perennial plants like citrus are largely unexplored due to long lifespans lengthening the time required to perform experiments, large field or greenhouse space required to grow tree crops, reduced access to diverse genotypes, and a lack of genomic resources.

There are multiple potential reasons why studies have observed variation in MAMP perception among related species. While some species may contain the same receptor homolog, the presence of the homolog does not always correlate with strong MAMP perception (Vetter *et al*. 2012, Trdá *et al*. 2013). In this study, the LYK5_*At*_ homologs in ‘Washington navel’ orange and ‘Tango’ mandarin are identical for allele 1 and only differ by two amino acid changes in allele 2. These polymorphisms have not been previously described as important for LYK5 receptor function (Cao *et al*. 2014). However, ‘Tango’ mandarin is a much stronger ROS responder to chitin than ‘Washington navel’ orange. Similarly, ‘Washington navel’ orange does not respond to flg22, while ‘Frost Lisbon’ lemon does. Similar to our LYK5 results, both responsive and nonresponsive genotypes contain a functional *FLS2* homolog when expressed in *Arabidopsis* or 1. *N. benthamiana*. In another study, Vetter *et al*. 2012 noted that the variation in FLS2 protein abundance for certain genotypes can reflect the variation in flg22 binding. While we were unable to determine protein abundance for LYK5 and FLS2 in citrus, both receptors were transcriptionally expressed at a similar basal level in responding and non-responding genotypes. Minor variations in the rice OsCERK1 co-receptor have been linked to variation in mycorrhizal symbiosis, thus it is possible minor allelic variation could also explain responsiveness to flagellin or chitin in citrus (Huang *et al*. 2020).

The segregation of immune response outputs has been observed previously, where flg22 from *C*Las induces defense gene induction but no ROS burst in ‘Sun Chu Sha Kat’ mandarin (Shi *et al*. 2018). Overexpression of the *N. benthamiana* FLS2 receptor in ‘Hamlin’ orange is able to confer flg22 responsiveness, indicating that boosting PRR expression may be a viable strategy to gain MAMP recognition (Hao *et al*. 2016). There is also a possibility that downstream signaling components may play a role in the presence and magnitude of immune responses. Signaling has not been investigated in detail in perennial crops, and further research may reveal if downstream signaling components have a role in altering MAMP responsiveness in citrus.

Flagellin perception is widespread amongst plants, mainly conferred by the receptor FLS2 (Saijo *et al*. 2018). Additional receptors were identified based on homology to *Arabidopsis* FLS2 in tomato, grapevine, citrus and rice (Robatzek *et al*. 2007; Trdá *et al*. 2013; Shi *et al*. 2016; Takai *et al*. 2008). From our study, LYK5 and CERK1 homologs are also widespread and cluster separately between dicots and monocots. Rice utilizes a different LysM receptor (CEBiP) along with the CERK1 co-receptor for chitin perception (Kaku *et al*. 2006, Lee *et al*. 2014, Shimizu *et al*. 2010). Cotton, a dicot, has a wall-associated kinase that interacts with LYK5 and CERK1 to promote chitin-induced dimerization (Wang *et al*. 2020). These results are consistent with an ancient acquisition of chitin perception in dicots, which may explain why a vast majority of the Rutaceae genotypes we evaluated are capable of producing an immune response to chitin. For MAMPs that can be recognized by a broad range of species, identifying receptors based on homology is a useful tactic.

Recent studies have identified immune receptor homologs that are capable of perceiving polymorphic flg22 epitopes that are not perceived by the canonical *Arabidopsis* FLS2 receptor. Flg22 from *Agrobacterium tumefaciens* contains several polymorphisms that prevent perception in *Arabidopsis* (Felix *et al*. 1999). Fürst *et al*. 2020 identified a novel flagellin-sensing receptor from wild grape with expanded ligand perception, FLS2^XL^, capable of sensing both canonical flg22 and the *Agrobacterium* flg22 epitopes. The *Ralstonia solanacearum* flg22 is also highly polymorphic and is not recognized by tomato (Pfund *et al*. 2004). Wei *et al*. 2020 identified a FLS2/BAK1 complex in soybean that is capable of sensing the *Ralstonia* flg22. In this study, we have identified Rutaceae genotypes that are capable of recognizing canonical csp22 and csp22_*C*Las_. In tomato, the CORE RLK is responsible for csp22 recognition (Wang *et al*. 2016). However, no obvious homolog of the *CORE* receptor has been identified in citrus genomes so far. While homology can be a fruitful approach to identify candidate receptors, convergent receptor evolution to recognize the same MAMP is also possible. There is likely an independently-evolved receptor that members of the Rutaceae possess to recognize csp22 epitopes. Comparative genomics of csp22-responsive and non-responsive citrus genotypes, or segregating populations if available, could be used to identify candidate receptor(s) for csp22 epitopes for future functional validation.

HLB induces some hallmarks of defense in susceptible plants, including callose and elevated ROS production, indicating that it can be a pathogen-triggered immune disease (Ma *et al*. 2022). The HLB susceptible orange genotypes we analyzed were unable to robustly respond to most MAMPs, unlike the more HLB tolerant ‘Frost Lisbon’ lemon. These results are consistent with weak, continuous, and ineffective defense activation in HLB susceptible citrus resulting in detrimental immune responses. It is possible that introduction of multiple PRRs capable of robustly inducing defense against *C*Las, including the csp22_*C*Las_ receptor, may result in active pathogen clearing. *C. ×aurantium* L. overexpressing the SA receptor and master regulator NPR1 exhibited increased tolerance to HLB and decreased pathogen titers (Dutt *et al*. 2015, Peng *et al*. 2021, Robertson *et al*. 2018). Appropriate regulation of defense and careful introduction of candidate receptors/genes should be considered with respect to HLB mitigation.

Our study highlights the diversity of immune response in a genetically diverse plant family. We identified relatives of citrus that are capable of responding to a wide variety of MAMPs, opening up new opportunities to study relatives with potential novel mechanisms of immune signaling. The transfer of novel receptors for MAMPs to susceptible plants can generate resistance to pathogens (Hao *et al*. 2016, Fürst *et al*. 2020, Wei *et al*. 2020). ROS-based immune phenotyping can be a high-throughput method to accelerate selection of promising individuals in a breeding program. Individuals that can respond to unique MAMPs are likely to have unique immune signaling components that can be transferred to susceptible citrus varieties. A greater understanding of immune perception repertoires in economically important plant genotypes will also facilitate the design stacks of receptors or signaling components for transfer and disease control. Similar strategies have resulted in durable resistance against fungal and oomycete pathogens in crop plants (Luo *et al*. 2021, Ghislain *et al*. 2018). This work has opened up interesting avenues to identify new receptors in non-model species and highlight genotypes capable of recognizing polymorphic pathogen features.

## Materials and Methods

### Plant materials and growth conditions

Eighty-six Rutaceae genotypes were tested for MAMP responsiveness under field and greenhouse conditions using mature trees (>5 years old). Field-grown trees in the Givaudan Citrus Variety Collection (GCVC) in Riverside, CA (http://www.citrusvariety.ucr.edu) were analyzed between the spring and fall from 2018-2021. Table S1 includes all genotypes analyzed, their accession IDs, and their location. When collecting Rutaceae samples, branches of selected Rutaceae genotypes were retrieved either from the greenhouse or the field, selecting branches with fully-expanded leaves that still retained flexibility. Branches were stored by placing the cut side of the branch into a wet floral block (Oasis^Ⓡ^ #10-00020-CASE) until processing.

*Arabidopsis thaliana* seeds (Col-0 or *lyk4/lyk5-2* mutant) were stratified for 2 days in the dark at 4°C before sowing onto soil or half-strength Murashige and Skoog (MS) media (Cao *et al*. 2014). Plants were grown in a Conviron growth chamber at 23°C and 70% relative humidity with a 10-h light/14-h dark photoperiod (100 μM m−2 s−1). 10- to 14-day-old seedlings grown on MS were used for MAPK phosphorylation and MTI marker gene induction assays, and four-week-old soil grown plants were used for ROS and MAPK assays.

*Nicotiana benthamiana* was grown in a growth chamber at 26°C with a 16-h light/8-h dark photoperiod (180 μM m−2 s−1). 30-day-old (before flowering) plants were used for *Agrobacterium*-mediated transient protein expression, and three- to 4-week-old plants were used to examine *LYK5* transgene expression.

### Microbe-associated molecular patterns (MAMPs)

Immunogenic epitopes for flg22 and csp22 peptides were synthesized using Genscript (≥ 95% purity, Piscataway, NJ, USA). Hexaacetyl-chitohexaose (chitin) (Megazyme #O-CHI6) was diluted in water. The canonical flg22 epitope (QRLSTGSRINSAKDDAAGLQIA) is based on the sequence information from *Pseudomonas aeruginosa*. The canonical csp22 sequence (AVGTVKWFNAEKGFGFITPDGG) is from *Pseudomonas syringae*. The csp22 *C*Las sequence (HRGSIKWYNPDKGYGFITPEGS) is identical to the sequence from the *C*Las strain psy62 (CLIBASIA_04060).

### Reactive oxygen species (ROS) burst assay

Leaf disks were collected using a #1 cork borer (4 mm) and floated overnight in 200 μL demineralized water in a Corning™ Costar™ 96-Well White Solid Plate (Fisher #07-200-589) with a plastic lid to prevent evaporation of the water. The subsequent day, water was replaced with 100 μL of an assay solution containing MAMP. The assay solution contained 20 μM L-012 (a luminol derivative from Wako Chemicals USA #120-04891), 10 mg mL−1 horseradish peroxidase (Sigma), and MAMP. Concentrations used for MAMP treatments were: 100 nM flg22, 10 μM chitin, 100 nM csp22, or 200 nM csp22_*C*Las_. Luminescence was measured using a GloMax®-Multi+ Reader (Promega) or TriStar LB 941 plate reader (Berthold Technologies). Significance of differences between experimental groups was determined using ANOVA and Tukey’s test, α = 0.05. At least 8 leaf discs were used in each replication, and the experiments were repeated at least three times. Samples that only responded once were considered variable.

For each plate, an average maximum RLUs for each tested MAMP-genotype combination were calculated from the maximum RLU of the 8 leaf disks after subtraction by the average water RLU. Across all ROS plates, average maximum RLUs were calculated based on all plates ran for each MAMP-genotype and a heatmap was created from the accumulated ROS data via custom R scripts (Github repository: DanielleMStevens/Divergent_citrus_response_to_PAMPs) including the following R packages: ComplexHeatmap (v2.5.1, Gu *et al*. 2016), circlize (v0.4.8, Gu *et al*. 2014).

### Mitogen-activated protein kinase (MAPK) induction assay

For MAPK induction assays in citrus, leaf disks were collected using a #6 cork borer (12 mm) from accessions grown in UC Davis or UC Riverside greenhouses and floated overnight in 1 mL deionized water in a 24-well tissue culture plate (VWR #10062-896) with a plastic lid to prevent evaporation of the water. The subsequent day, water was replaced with 500 μL of either water or water containing MAMP before pressure-infiltrating for 2 minutes at 30 mm Hg in a vacuum desiccator (SP Bel-Art #F42025-0000). Leaf disks were collected at 0, 10, and 20 minutes after vacuum infiltration, flash frozen in liquid nitrogen, and ground up with pestles attached to an electric grinder (Conos AC-18S electric torque screwdriver) before adding 200 μL extraction buffer and grinding until homogenous. Protein extraction buffer contained 50 mM HEPES (pH 7.5), 50 mM NaCl, 10 mM EDTA, 0.2% Triton X-100, Pierce™ Protease Inhibitor Mini Tablets, EDTA-free (Thermo #A32955), Pierce™ Phosphatase Inhibitor Mini Tablets (Thermo #A32957). Samples were centrifuged at 15,000 rpm for 10 minutes to pellet cell debris.

Protein concentrations were quantified with the Pierce 660 nm Protein Assay Reagent (Thermo #22660) with Ionic Detergent Compatibility Reagent (Thermo #22663). MAPKs were visualized by anti-p44/42 MPK immunoblotting (1:2000, Cell Signaling Technology #4370L) with goat anti-rabbit HRP secondary antibody (1:3000, Bio-Rad #170-5046). Membranes were developed using the SuperSignal West Pico Chemiluminescent Substrate kit (Fisher #PI34578) and visualized on a ChemiDoc™ Touch Gel Imaging System (BioRad #1708370).

For MAPK induction assays in *Arabidopsis*, 5 day-old seedlings were transplanted into an 48-well tissue culture plate (Costar #3548) supplemented with half-strength MS liquid media. After 9 days, MS liquid media were replaced with 500 μL of either water or water containing 10 μM chitin. Three seedlings per treatment were collected at 0 and 15 minutes after chitin induction. Protein extraction and western blotting were conducted as described above.

### qPCR

To examine the expression of *LYK5* homologs, citrus leaves were harvested to make leaf punches with a #9 cork borer (22.5 mm). Each leaf disk was placed in a 12-well plate (VWR #10062-894) with 1 mL water and kept overnight at room temperature to allow the samples to recover from wounding. The subsequent day, water was replaced with either 1 mL water or water containing 1 mL 10 μM chitin before vacuum-infiltrating for 1.5 minutes at 30 mm Hg in a vacuum desiccator (SP Bel-Art #F42025-0000). Three leaf disks per treatment were collected at 0 and 24 h post infiltration and RNA extraction was performed as described above. To examine the expression of *FLS2* homologs, citrus leaves were syringe-infiltrated with water or 1μM flg22 and plant samples harvested 6hpi. RNA extraction was performed as described above. Samples for resting state expression of *LYK5* and *FLS2* homologs were taken at 0h as described above without infiltrating with water or MAMP.

To examine the expression of MTI marker genes, citrus leaves were infiltrated on the tree with either water or 10 μM MAMP. At 0 and 24 h after infiltration, a #6 cork borer (12 mm) was used to make 6 leaf punches of infiltrated areas per sample. Leaf punches were manually ground with liquid nitrogen into a fine powder before transferring the powder to tubes to perform RNA extractions. RNA was extracted from plant samples with TRIzol (Fisher #15596018), following the manufacturer’s instructions. DNase treatments for RNA preps were performed with RQ1 RNase-Free DNase (Promega #PR-M6101). cDNA synthesis was performed with the MMLV Reverse Transcriptase (Promega #PRM1705) kit.

A table of qPCR primers can be found in Table S2. Citrus *GAPDH* (glyceraldehyde-3-phosphate dehydrogenase, Pang *et al*. 2020) was used as the reference gene for qRT-PCR reactions. qPCR reactions were performed with SsoFast EvaGreen Supermix with Low ROX (BioRad #1725211) in a 96-well white PCR plate (BioRad #HSP9601) according to the manufacturer’s instructions. Fold-induction of gene expression determined using the 2^−ΔΔCt^ method (Livak and Schmittgen 2001), normalizing to water-treated and 0-hour timepoints.

### Phylogenetic analyses and receptor comparisons

Plant genotypes used to build LYK5 and CERK1 phylogenies can be found in Table S3 and S4, respectively. Using the *Arabidopsis thaliana* (NCBI taxid: 3702) CERK1_*At*_ (NCBI RefSeq: NP_566689.2) and LYK5_*At*_ (NCBI RefSeq: NP_180916.1) as queries, a blastp search was performed to mine for homologs across higher plant species (BLAST Suite v2.11.0+, query coverage cutoff 80%, E-value cutoff 1e-50, defaults otherwise). Partial protein hits were removed. Hits were then compared to all LysM RLKs in *A. thaliana* (CERK1, LYK2, LYK3, LYK4, and LYK5) and homologs closer to another LYK were removed. LYK5_*At*_ homologs from members of the Solanaceae family were removed because they were closer to LYK4_*At*_ than LYK5_*At*_. A multiple sequence alignment of receptor homologs was built using MAFFT (v7.310, -- reorder, --maxiterate 1000, --localpair, defaults otherwise). TrimAl was used to trim each multiple sequence alignment for large gaps (v1.4.rev15, automated1, defaults otherwise). A maximum likelihood tree was built from the alignment using iqtree (v2.1.2, —bb 1000, -T AUTO, -st AA, -v -m MFP -safe, defaults otherwise), mid-rooted, and visualized using R packages phangorn (v2.7.1) and ggtree (v3.1.2.991).

We sought to characterize the number of chitin-binding LysM domains in LYK5 and CERK1 homologs. The standard domain prediction software Interproscan was unable to accurately predict even well characterized LYK5_*At*_ and CERK1_*At*_ receptors. To improve assessment of LysM domain frequency, we used homology and a hidden Markov model approach by blastp and HMMER, respectively. Hmmersearch using the LysM domain (query ID: PF01476.19) as a query identified the number of LysM domains (hmmer v3.1b2, -E 1e-5, defaults otherwise) (Eddy 2011). Manually extracted LysM domains from *A. thaliana* Col-0 were used to build a local blast database to calculate similarity of each LysM domain (coverage cutoff 80%, E-value cutoff 1e-50, defaults otherwise). All LysM domain analyses were plotted onto the receptor trees in R.

Ectodomains of all receptor homologs were extracted and an all-by-all comparison was computed using blastp. Similarity to either CERK1_*At*_ and LYK5_*At*_ or all citrus homologs to each other was plotted using R packages ggplot2 (v3.3.5) and ggbeeswarm (v0.6.0). Weblogos were generated from the multiple sequence alignment corresponding to LYK5_*At*_ residues Y128 and S206 using WebLogo3 (Crooks *et al*. 2004).

The phylogenetic tree of citrus CORE homologs was built with CORE homologs from other solanaceous plants (*Solanum lycopersicum*: Solyc03g096190; *Solanum pennellii*: XP_015068909.1; *Nicotiana benthamiana*: Niben101Scf02323g01010.1; *Nicotiana sylvestris*: XP_009803840.1; *Nicotiana tabacum*: XP_016470062.1). Protein sequences were aligned via MAFFT, tree building was performed by iqtree, and were visualized in R similarly as described above. Sequences of cloned CORE homologs from ‘Frost nucellar Eureka’ lemon and ‘Washington navel’ orange are deposited in GenBank (ON863917 and ON863918, respectively).

For more information and raw files see Github repository: DanielleMStevens/Divergent_citrus_response_to_PAMPs.

### Cloning citrus LYK5 homologous sequences and transcomplementation in Arabidopsis

Putative LYK5 sequences from Australian finger lime, ‘Rio Red’ grapefruit, ‘Frost Lisbon’ lemon, ‘Tango’ mandarin and Eremolemon were amplified using iProof DNA polymerase (Bio-Rad #BR0114). The PCR products were initially cloned into a pENTR™/D-TOPO™ backbone (Invitrogen #K2400-20), then *LYK5* from ‘Tango’ mandarin (TM) was selected and inserted into a modified pGWB14 binary destination vector with the *Arabidopsis* ubiquitin 10 promoter using Gateway LR Clonase™ II enzyme mix (Invitrogen #11791-100). The *Arabidopsis lyk4/lyk5-2* mutant (Cao *et al*. 2014) was transformed using a floral dip method with pUBQ10::TM_LYK5-HA and pUBQ10::LYK5-HA from *Arabidopsis* (Zhang et al. 2006). Experiments were performed with T4 homozygous lines.

### Virus-induced gene silencing (VIGS)

VIGS is performed as described in (Chakravarthy *et al*. 2010). Two-week old *Nicotiana benthamiana* seedlings were infiltrated with pTRV1(RNA1), along with silencing constructs: pTRV2:*GUS*, pTRV2:*PDS*, pTRV2:*FLS2* (Chakravarthy *et al*. 2010). After two to three weeks, silenced plants were infiltrated with *Agrobacterium* harboring FLS2 constructs for transient expression and ROS assays. *Arabidopsis thaliana* Col-0 *FLS2*, ‘Washington navel’ orange *FLS2* and ‘Frost Lisbon’ lemon *FLS2* homologs were amplified and inserted into a modified pGWB14 binary destination vector with the *Arabidopsis* ubiquitin 10 promoter as described above. Plasmids were transformed via electroporation into *Agrobacterium* C58C1 and VIGS silenced *Nicotiana benthamiana* plants were infiltrated with *Agrobacterium* suspensions of OD_600_ 0.6. Forty-eight hours after infiltration, leaf disks were collected using a #1 cork borer (4 mm) for ROS assays after challenging with 100 nM flg22 as described above. To visualize the expression of FLS2 homologs, additional leaf disks were collected using a #7 cork borer at 48 hpi for protein extraction. Leaf disks were homogenized in 100 μL Laemmli buffer and boiled for 5 min. Western blotting was conducted as described above and visualized with anti HA-HRP antibody (Roche 39 #12013819001; 1:2,000).

### *Transient expression of CORE in* Nicotiana

The closest CORE homologs from ‘Frost nucellar Eureka’ lemon and ‘Washington navel’ orange were amplified using iProof DNA polymerase (Bio-Rad #BR0114). PCR products were cloned into a pEARLY103 backbone (Earley *et al*. 2006) with expression mediated by a 35S promoter, and plasmids were transformed via electroporation into *Agrobacterium* GV3101. *Nicotiana benthamiana* CORE was used as a positive control (Wang *et al*. 2016). To test the function of CORE homologs, young (non-flowering) *Nicotiana benthamiana* plants were infiltrated with *Agrobacterium* suspensions of OD_600_ 0.25. Twenty-four hours after infiltration, leaf disks were collected using a #1 cork borer (4 mm) for ROS Assays as described above. To visualize the expression of CORE homologs, additional leaf disks were collected using a #7 cork borer at 48 hpi for protein extraction. Leaf disks were homogenized in 100 μL Laemmli buffer and boiled for 5 min. Western blotting was conducted as described above and visualized with anti GFP-HRP antibody (Miltenyi Biotec #130-091-833).

### *Comparison of* LYK5 and *FLS2 homologs in citrus*

Genome sequences surrounding *LYK5* and *FLS2* homologs in cultivated citrus were compared based on BLASTp hits with an e-value of 0.001 (Altschul, et al. 1990) against *de novo* assembled genomes. Contigs were assembled using wtgbt2 (Ruan and Li *et al*. 2020) and long-read sequencing was performed on the Pacific Biosciences Sequel II platform in the CLR sequencing mode. LYK5 and FLS2 protein sequences were aligned using MUSCLE v5.1 (Edgar 2021).

## Acknowledgements and Funding

We thank Dr. Melanie Schori (National Germplasm resources Laboratory Botanist, USDA-ARS) for help with the taxonomy of Rutaceae and the Citrus Research Board for their continued support (Project 5200-171 to DKS and 5200-157 to GC). We thank Mikeal Roose for funding the ‘Tango’ mandarin genome assembly and Georg Felix for providing a binary vector containing *Nicotiana CORE* and Erika Roxana Espinoza for assisting with ROS assays. Funding was provided by USDA-NIFA grant no. 2019-70016-29796 and NIH grant no. R35GM136402 awarded to GC. DMS was supported by the USDA-NIFA predoctoral fellowship grant no. 2021-67034-35049. TL was supported by the China Scholarship Council No. 201906300032.

## Author Contributions

JF, TT, CR, and GC designed the research, and JT, TL, JF, TT, CR, RP, and JW performed the research. JT, TL, DMS, JF, TT, ST, CR and GC analyzed the data, and JT, TL, DMS, and GC wrote the paper.

DKS and EAD provided citrus sequencing data.

TLK provided guidance and access to greenhouse-grown Rutaceae genotypes.

## Data Availability

The data that support the findings of this study are available, with raw data available from the corresponding author upon request. The sequences of contigs containing *LYK5* from ‘Washington navel’ orange and ‘Tango’ mandarin are deposited in GenBank (*Citrus reticulata* ON685188, *Citrus ×aurantium* L. ON685189 and *Citrus reticulata* ON685190, *Citrus ×aurantium* L. ON685191). The sequences of contigs containing *FLS2* from ‘Washington navel’ orange and ‘Frost Lisbon’ lemon are deposited in Genbank (OP718785, OP718786, OP718788, OP718789). Software used to build the heatmap and parts of Figure 5 are deposited in GitHub under the repository DanielleMStevens/Divergent_citrus_response_to_PAMPs. Sequences of cloned CORE homologs from ‘Frost nucellar Eureka’ lemon and ‘Washington navel’ orange in **Supplemental Information Figure 8B** are deposited in GenBank (ON863917 and ON863918, respectively).

## Supplemental Information

**Supplemental Information Table S1** contains information on the Rutaceae genotypes used for this study.

**Supplemental Information Table S2** contains information on primers used for this study.

**Supplemental Information Table S3** contains information on the LYK5 sequences used to build Figure 5a.

**Supplemental Information Table S4** contains information on the CERK1 sequences used to build Figure 5a.

---

**Supplemental Information Figure 1.**
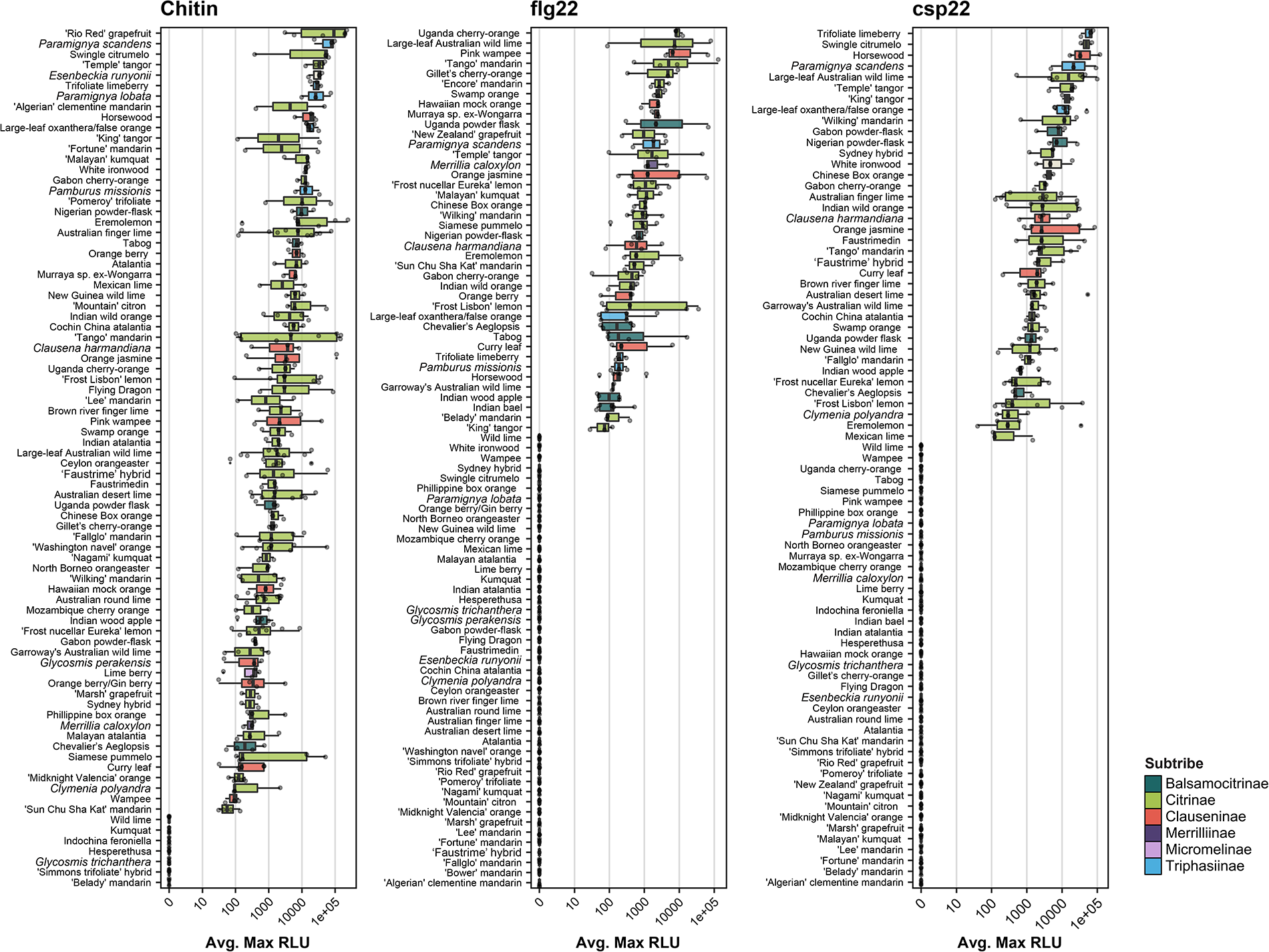
Box plots containing the distribution of ROS data from Figure 1 for all non-variable Rutaceae genotypes. Data points on box plots represent the average max RLU for an individual experiment, with n = 8 leaf disks per experiment. M*ax RLU = (max RLU MAMP – max RLU water)*. <90 RLU is the threshold for no response. The max RLU are plotted on a log10 scale. The bar within the box plot depicts the median of the data, where the box boundaries represent the interquartile range (between the 25th and 75th percentiles) of the data. Box whiskers represent the minimum or maximum values of the data within 1.5× of the interquartile range.

**Supplemental Information Figure 2.**
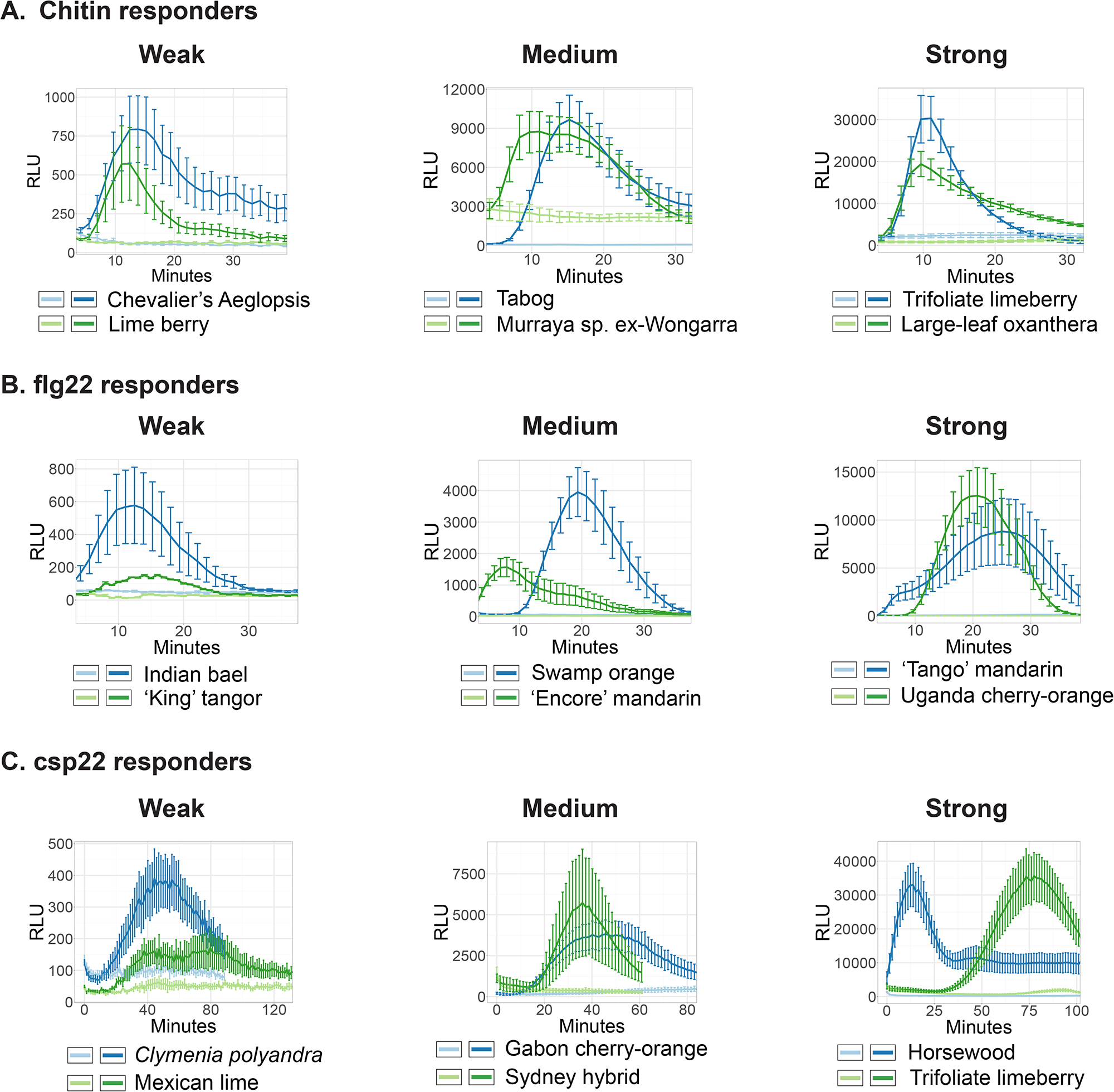
Representative ROS curves for the genotypes shown in Figure 2. Light-colored lines represent water treatments, dark-colored lines represent MAMP treatments. ROS curve dynamics for weak (left), medium (middle), and strong (right) responders to MAMPs in Figure 2. The MAMPs used are canonical features in the following concentrations: chitin (A, 10 μM), flg22 (B, 100 nM), csp22 (C, 100 nM). Water treatment was used as a negative control. Criteria for the response categories: “strong” responders are in the top 25th percentile, “medium” responders are between the 25th and 75th percentiles, and “weak” responders are in the bottom 25th percentile. RLU = relative light units. Error bars represent the standard error of the means (SEM) of n = 8 leaf disks per experiment.

**Supplemental Information Figure 3.**
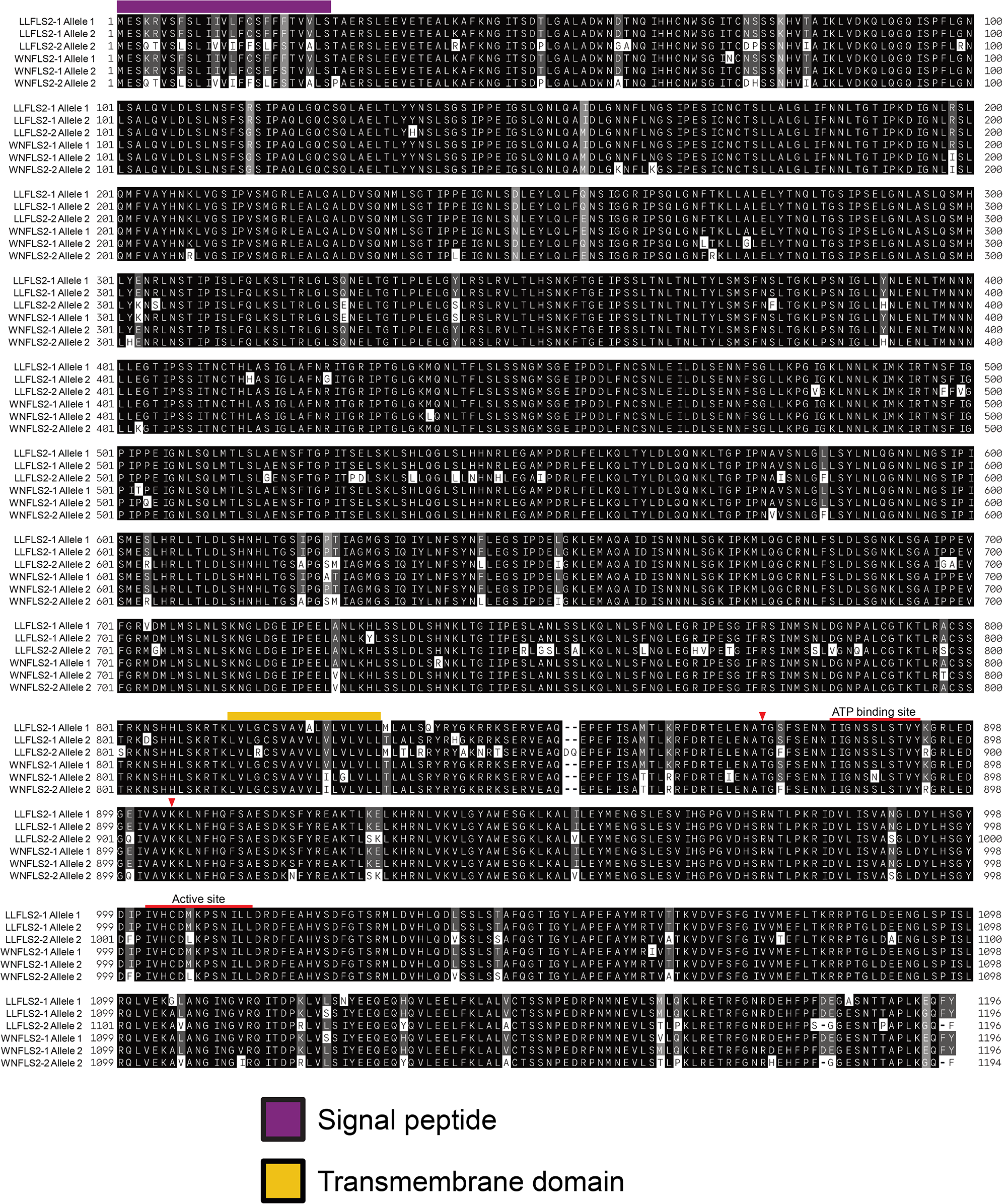
Amino acid alignments of FLS2 homologs. Sequences are color-coded according to functional domain. Red arrows indicate residues crucial for flg22 binding or kinase activity in *Arabidopsis*. LL = ‘Frost Lisbon’ lemon, WN = ‘Washington navel’ orange. Allele 1 refers to haplotype 1 and allele 2 refers to haplotype 2. The NCBI accession numbers for ‘Frost Lisbon’ lemon FLS2-1 Allele 1 is OP718788; FLS2-1 Allele 2 is OQ572427; FLS2-2 Allele 2 is OP718789. The ‘Washington navel’ orange FLS2-1 Allele 1 is OP718785; FLS2-1 Allele 2 is OQ572428; FLS2-2 Allele 2 is OP718786. The ‘Washington navel’ orange FLS2-2 Allele 1 (haplotype 1) is a pseudogene due to a premature stop codon, thus not shown in the alignment, its NCBI accession number is OQ572429.

**Supplemental Information Figure 4.**
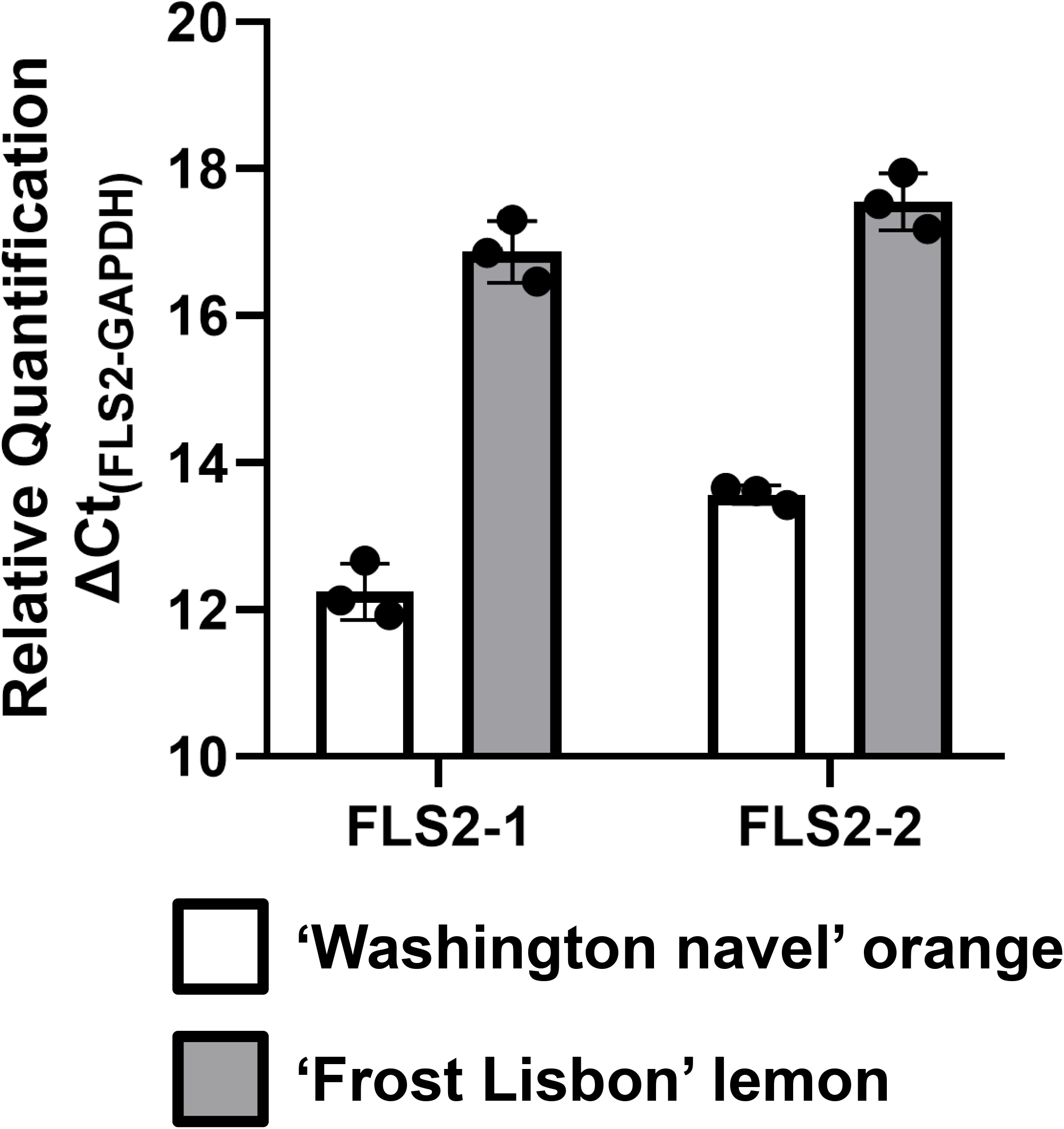
Baseline expression of *FLS2-1* and *FLS2-2*. ΔCt of the citrus *FLS2-1* and *FLS2-2* transcripts measured via qPCR at resting state, using citrus *GAPDH* as a reference gene. Error bars represent standard deviation (n = 3 biological replicates). A higher ΔCt indicates lower transcript expression.

**Supplemental Information Figure 5.**
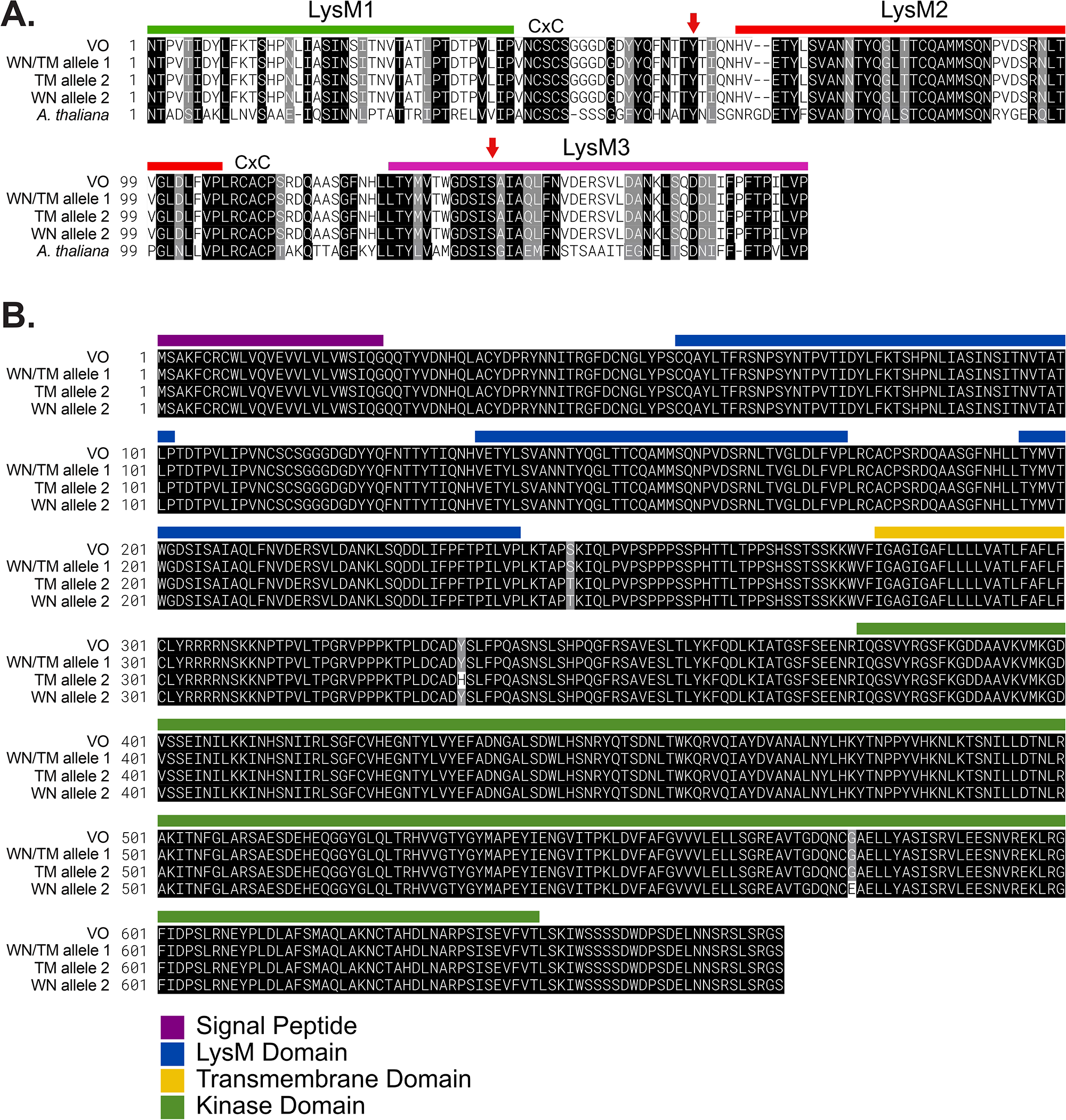
Amino acid alignments of LYK5 homologs. A. Alignment of the LYK5 LysM domains between citrus and *Arabidopsis thaliana*, with CXC motifs that separate the LysM domains indicated. Colors represent the different LysM domains. Red arrows indicate residues important for chitin binding in Arabidopsis (Y128 and S206, Cao *et al*. 2014). VO = ‘Midknight Valencia’ orange, WN = ‘Washington navel’ orange, TM = ‘Tango’ mandarin. The WN and TM allele 1 are identical. B. Alignment of full length citrus LYK5 homologs, color-coded according to functional domain. The NCBI accession numbers for ‘Midnight Valencia’ orange is XP_006494433.1. The allele 1 of ‘Washington navel’ orange and ‘Tango’ mandarin is identical, with the sequence deposited as ON685190. The ‘Washington navel’ orange allele 2 is ON685191, and the ‘Tango’ mandarin allele 2 is OP718787.

**Supplemental Information Figure 6.**
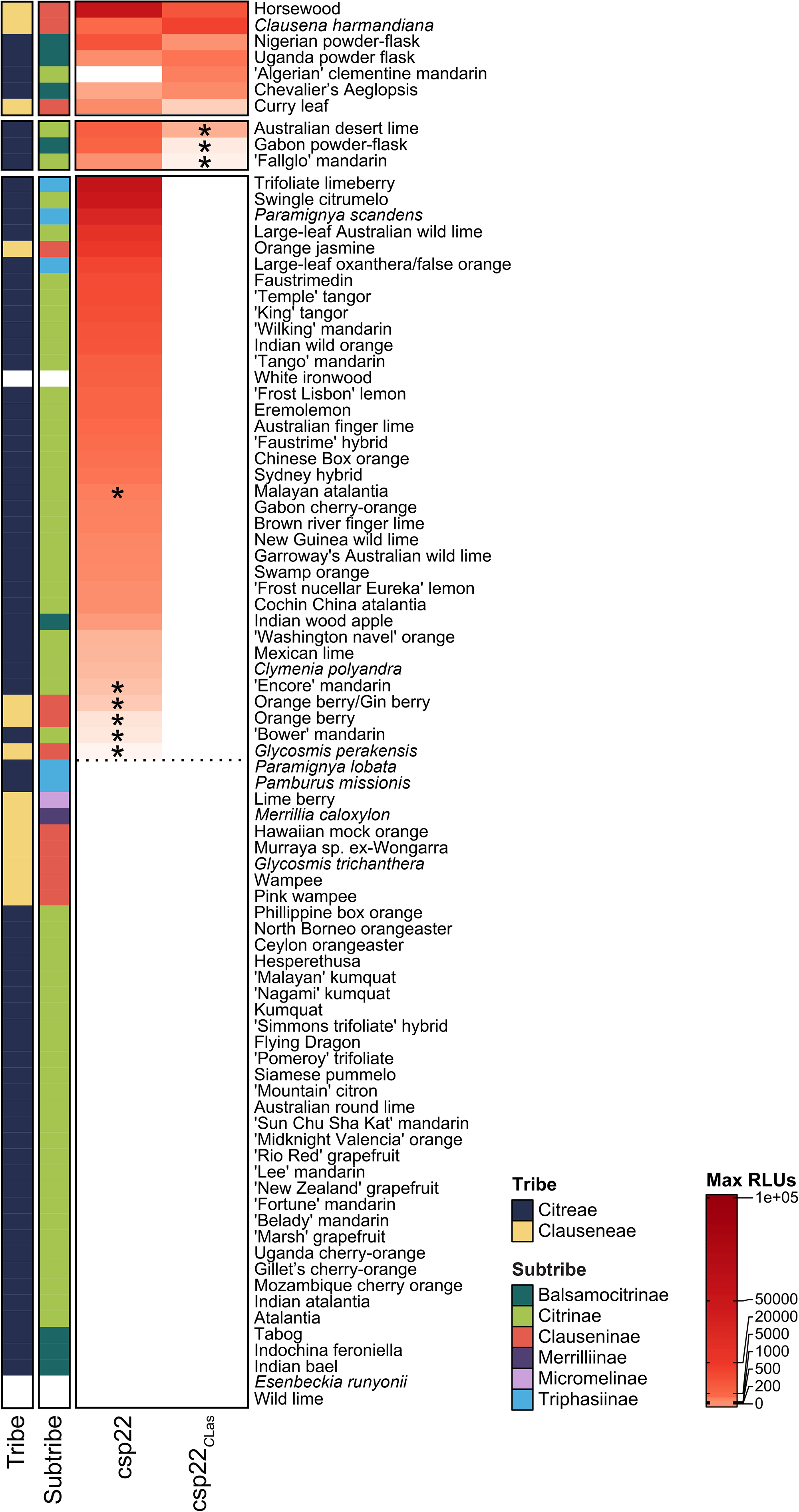
Full heat map featuring all genotypes tested with either canonical csp22 or *Candidatus* Liberibacter asiaticus csp22, related to Figure 7. Heat map compiling average max relative light units (RLUs) from ROS assays. Max RLUs are averages of at least three independent experiments and are represented as a heatmap, where *max RLU = (max RLU MAMP – max RLU water)*. <90 RLU is the threshold for no response. Asterisks indicate genotypes that exhibit a variable response, where one or two independent experiments revealed a response. The MAMPs used for treatments are canonical csp22 and csp22 from *Candidatus* Liberibacter asiaticus (csp22_*C*Las_).

**Supplemental Information Figure 7.**
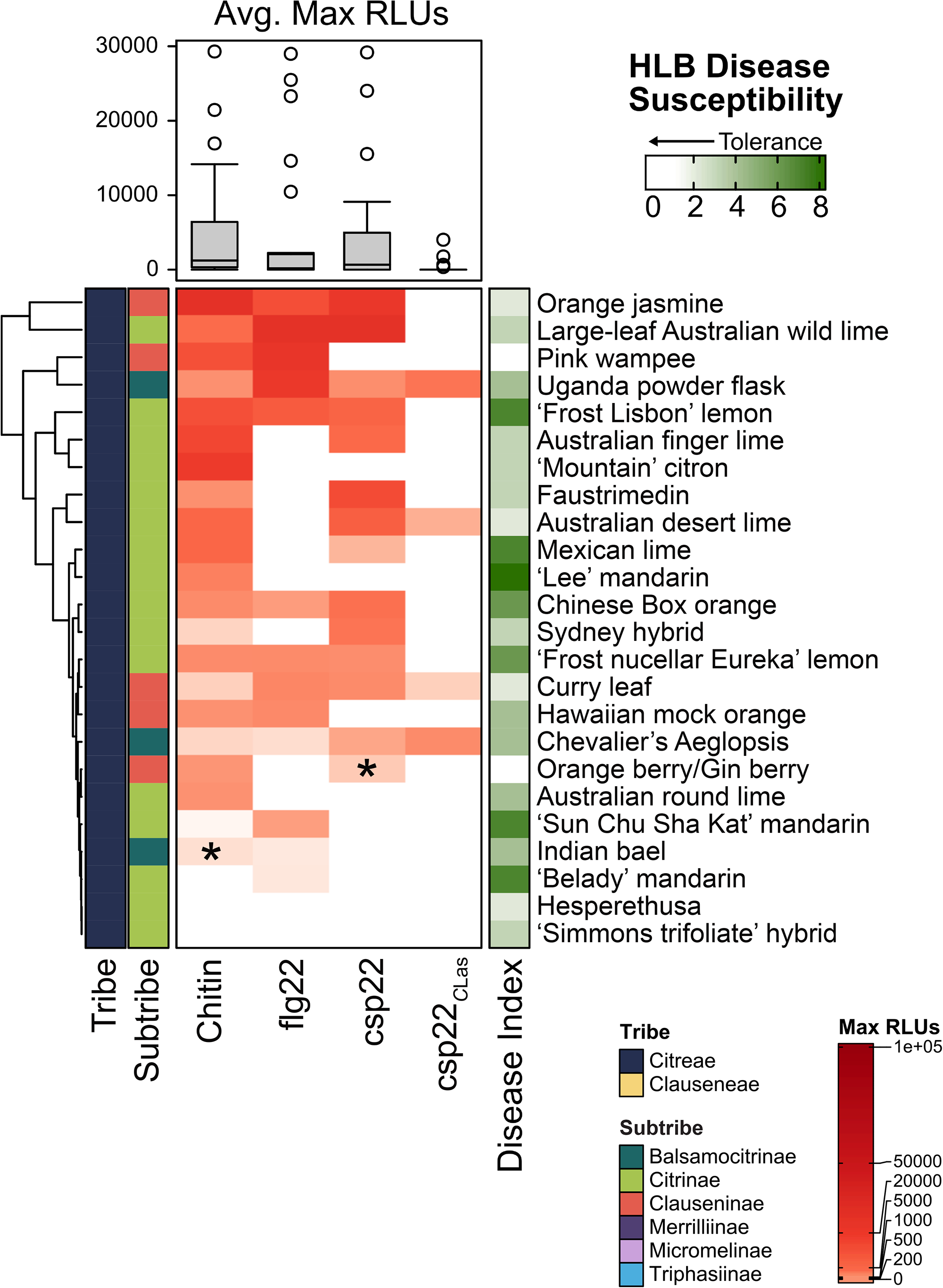
Visualization of MAMP perception with HLB disease susceptibility. Heat map compiling average max relative light units (RLUs) from reactive oxygen species (ROS) assays in genotypes within the Rutaceae family, organized by MAMP and phylogenetic relationship. Max RLUs are averages of at least three independent experiments and are represented as a heatmap, where *max RLU = (max RLU MAMP – max RLU water)*. <90 RLU is the threshold for no response. Asterisks indicate genotypes that exhibit a variable response, where one or two independent experiments revealed a response. The MAMPs used are canonical features in the following concentrations: chitin (10 μM), flg22 (100 nM), csp22 (100 μM), as well as the csp22 from *Candidatus* Liberibacter asiaticus, the bacterium associated with citrus Huanglongbing (200 nM). Asterisks indicate genotypes that exhibit a variable response, where one or two independent experiments revealed a response. HLB disease susceptibility is rated on a 1-8 scale, with 0 representing no *C*Las multiplication and 8 representing severe disease symptoms and bacterial titers from Ramadugu *et al*. 2016.

**Supplemental Information Figure 8.**
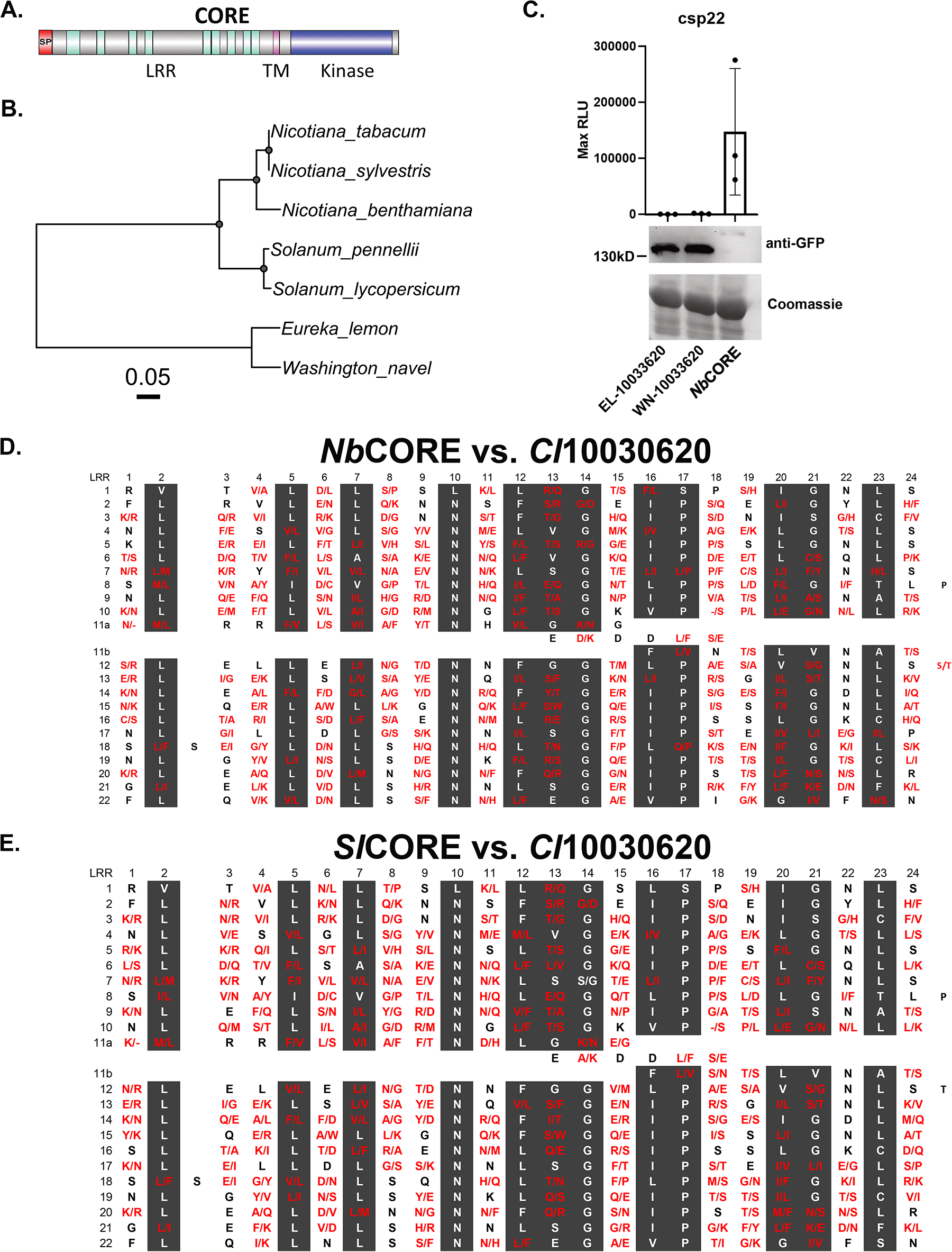
The closest citrus CORE homologs from ‘Washington navel’ orange (WN) and ‘Frost nucellar’ Eureka lemon (EL) cannot perceive csp22. A. Domain architecture of CORE highlighting four features: SP= signal peptide, LRR= leucine-rich repeats, TM= transmembrane domain and kinase domain. B. Mid-rooted phylogeny of CORE from *Nicotiana* and *Solanum* species (*S. lycopersicum*: Solyc03g096190; *S. pennellii*: XP_015068909.1; *N. benthamiana*: Niben101Scf02323g01010.1; *N. sylvestris*: XP_009803840.1; *N. tabacum*: XP_016470062.1), and their closest homologs from EL and WN (based on *Citrus clementina* Ciclev10030620m homolog). 1000 ultrafast bootstrap replicates were calculated and values over 90 were plotted as a gray dot. C. The closest citrus CORE homologs from LL (*Citrus ×limon (L.)* Osbeck) and WN (*Citrus ×aurantium* L.) are unable to produce ROS in *Nicotiana benthamiana* after csp22 treatment. The positive control, CORE from *Nicotiana benthamiana* (*Nb*), is able to induce ROS production in response to csp22. Bottom: anti-GFP immunoblot analyses demonstrating construct expression. *Nb* CORE runs at a higher molecular weight, likely due to glycosylation. D. Amino acid comparison of 22 LRRs (xLxxLxLxxNxLS/TGxIPxxLGxLx consensus) from *Nicotiana benthamiana* (*Nb*) CORE and EL (Ciclev10030620m). In red are dissimilar amino acids. E. Amino acid comparison of 22 LRRs (xLxxLxLxxNxLS/TGxIPxxLGxLx consensus) from tomato (*Sl*) CORE and EL (Ciclev10030620m).

## References

1. Altschul, S.F., Gish, W., Miller, W., Myers, E.W., Lipman, D.J., 1990. Basic local alignment search tool. Journal of Molecular Biology 215, 403–410. https://doi.org/10.1016/S0022-2836(05)80360-2

2. Alves, M.N., Lopes, S.A., Raiol-Junior, L.L., Wulff, N.A., Girardi, E.A., Ollitrault, P., Peña, L., 2021. Resistance to ‘Candidatus Liberibacter asiaticus,’ the Huanglongbing Associated Bacterium, in Sexually and/or Graft-Compatible Citrus Relatives. Frontiers in Plant Science 11.

3. Antolín-Llovera, M., Petutsching, E.K., Ried, M.K., Lipka, V., Nürnberger, T., Robatzek, S., Parniske, M., 2014. Knowing your friends and foes – plant receptor-like kinases as initiators of symbiosis or defence. New Phytologist 204, 791–802. https://doi.org/10.1111/nph.13117

4. Appelhans, M.S., Bayly, M.J., Heslewood, M.M., Groppo, M., Verboom, G.A., Forster, P.I., Kallunki, J.A., Duretto, M.F., 2021. A new subfamily classification of the Citrus family (Rutaceae) based on six nuclear and plastid markers. TAXON 70, 1035–1061. https://doi.org/10.1002/tax.12543

5. Asai, T., Tena, G., Plotnikova, J., Willmann, M.R., Chiu, W.-L., Gomez-Gomez, L., Boller, T., Ausubel, F.M., Sheen, J., 2002. MAP kinase signalling cascade in Arabidopsis innate immunity. Nature 415, 977–983. https://doi.org/10.1038/415977a

6. Bigeard, J., Colcombet, J., Hirt, H., 2015. Signaling Mechanisms in Pattern-Triggered Immunity (PTI). Molecular Plant, Cell Signaling 8, 521–539. https://doi.org/10.1016/j.molp.2014.12.022

7. Bové, J.M., 2006. HUANGLONGBING: A DESTRUCTIVE, NEWLY-EMERGING, CENTURY-OLD DISEASE OF CITRUS. Journal of Plant Pathology 88, 7–37.

8. Cao, Y., Liang, Y., Tanaka, K., Nguyen, C.T., Jedrzejczak, R.P., Joachimiak, A., Stacey, G., 2014. The kinase LYK5 is a major chitin receptor in Arabidopsis and forms a chitin-induced complex with related kinase CERK1. eLife 3, e03766. https://doi.org/10.7554/eLife.03766

9. Caruso, M., Smith, M.W., Froelicher, Y., Russo, G., Gmitter, F.G., 2020. Chapter 7 - Traditional breeding, in: Talon, M., Caruso, M., Gmitter, Fred G. (Eds.), The Genus Citrus. Woodhead Publishing, pp. 129–148. https://doi.org/10.1016/B978-0-12-812163-4.00007-3

10. Castle, W.S., 2010. A Career Perspective on Citrus Rootstocks, Their Development, and Commercialization. HortScience 45, 11–15. https://doi.org/10.21273/HORTSCI.45.1.11

11. Chen, Y., Bendix, C., Lewis, J.D., 2019. Comparative genomics screen identifies microbe-associated molecular patterns from Candidatus Liberibacter sp. that elicit immune responses in plants. Mol. Plant. Microbe. Interact. https://doi.org/10.1094/MPMI-11-19-0309-R

12. Cheng, J.H.T., Bredow, M., Monaghan, J., diCenzo, G.C., 2021. Proteobacteria Contain Diverse flg22 Epitopes That Elicit Varying Immune Responses in Arabidopsis thaliana. MPMI 34, 504–510. https://doi.org/10.1094/MPMI-11-20-0314-SC

13. Cifuentes-Arenas, J.C., Beattie, G.A.C., Peña, L., Lopes, S.A., 2019. Murraya paniculata and Swinglea glutinosa as Short-Term Transient Hosts of ‘Candidatus Liberibacter asiaticus’ and Implications for the Spread of Huanglongbing. Phytopathology® 109, 2064–2073. https://doi.org/10.1094/PHYTO-06-19-0216-R

14. Coletta-Filho, H.D., Castillo, A.I., Laranjeira, F.F., de Andrade, E.C., Silva, N.T., de Souza, A.A., Bossi, M.E., Almeida, R.P.P., Lopes, J.R.S., 2020. Citrus Variegated Chlorosis: an Overview of 30 Years of Research and Disease Management. Trop. plant pathol. 45, 175–191. https://doi.org/10.1007/s40858-020-00358-5

15. Couto, D., Zipfel, C., 2016. Regulation of pattern recognition receptor signalling in plants. Nat Rev Immunol 16, 537–552. https://doi.org/10.1038/nri.2016.77

16. Crooks, G.E., Hon, G., Chandonia, J.-M., Brenner, S.E., 2004. WebLogo: A Sequence Logo Generator. Genome Res. 14, 1188–1190. https://doi.org/10.1101/gr.849004

17. Dalio, R.J.D., Magalhães, D.M., Rodrigues, C.M., Arena, G.D., Oliveira, T.S., Souza-Neto, R.R., Picchi, S.C., Martins, P.M.M., Santos, P.J.C., Maximo, H.J., Pacheco, I.S., De Souza, A.A., Machado, M.A., 2017. PAMPs, PRRs, effectors and R-genes associated with citrus–pathogen interactions. Ann Bot 119, 749–774. https://doi.org/10.1093/aob/mcw238

18. Das, A.K., 2003. Citrus canker-A review. Journal of Applied Horticulture 5, 52–60.

19. Dutt, M., Barthe, G., Irey, M., Grosser, J., 2015. Transgenic Citrus Expressing an Arabidopsis NPR1 Gene Exhibit Enhanced Resistance against Huanglongbing (HLB; Citrus Greening). PLOS ONE 10, e0137134. https://doi.org/10.1371/journal.pone.0137134

20. Earley, K.W., Haag, J.R., Pontes, O., Opper, K., Juehne, T., Song, K., Pikaard, C.S., 2006. Gateway-compatible vectors for plant functional genomics and proteomics. The Plant Journal 45, 616–629. https://doi.org/10.1111/j.1365-313X.2005.02617.x

21. Eddy, S.R., 2011. Accelerated Profile HMM Searches. PLOS Computational Biology 7, e1002195. https://doi.org/10.1371/journal.pcbi.1002195

22. Edgar, R.C., 2021. MUSCLE v5 enables improved estimates of phylogenetic tree confidence by ensemble bootstrapping. https://doi.org/10.1101/2021.06.20.449169

23. Erwig, J., Ghareeb, H., Kopischke, M., Hacke, R., Matei, A., Petutschnig, E., Lipka, V., 2017. Chitin-induced and CHITIN ELICITOR RECEPTOR KINASE1 (CERK1) phosphorylation-dependent endocytosis of Arabidopsis thaliana LYSIN MOTIF-CONTAINING RECEPTOR-LIKE KINASE5 (LYK5). New Phytologist 215, 382–396. https://doi.org/10.1111/nph.14592

24. Ference, C.M., Gochez, A.M., Behlau, F., Wang, N., Graham, J.H., Jones, J.B., 2018. Recent advances in the understanding of Xanthomonas citri ssp. citri pathogenesis and citrus canker disease management. Mol Plant Pathol 19, 1302–1318. https://doi.org/10.1111/mpp.12638

25. Fürst, U., Zeng, Y., Albert, M., Witte, A.K., Fliegmann, J., Felix, G., 2020. Perception of Agrobacterium tumefaciens flagellin by FLS2XL confers resistance to crown gall disease. Nat. Plants 6, 22–27. https://doi.org/10.1038/s41477-019-0578-6

26. Gmitter, F.G., Hu, X., 1990. The possible role of Yunnan, China, in the origin of contemporary citrus species (rutaceae). Econ Bot 44, 267–277. https://doi.org/10.1007/BF02860491

27. Gómez-Gómez, L., Boller, T., 2000. FLS2: An LRR Receptor–like Kinase Involved in the Perception of the Bacterial Elicitor Flagellin in Arabidopsis. Molecular Cell 5, 1003– 1011. https://doi.org/10.1016/S1097-2765(00)80265-8

28. Gu, Z., Eils, R., Schlesner, M., 2016. Complex heatmaps reveal patterns and correlations in multidimensional genomic data. Bioinformatics 32, 2847–2849. https://doi.org/10.1093/bioinformatics/btw313

29. Gu, Z., Gu, L., Eils, R., Schlesner, M., Brors, B., 2014. circlize implements and enhances circular visualization in R. Bioinformatics 30, 2811–2812. https://doi.org/10.1093/bioinformatics/btu393

30. Hao, G., Pitino, M., Duan, Y., Stover, E., 2016. Reduced Susceptibility to Xanthomonas citri in Transgenic Citrus Expressing the FLS2 Receptor From Nicotiana benthamiana. Molecular Plant-Microbe Interactions. https://doi.org/10.1094/MPMI-09-15-0211-R

31. Hodges, A.W., Spreen, T.H., n.d. Economic Impacts of Citrus Greening (HLB) in Florida, 6.

32. Huang, R., Li, Z., Mao, C., Zhang, H., Sun, Z., Li, H., Huang, C., Feng, Y., Shen, X., Bucher, M., Zhang, Z., Lin, Y., Cao, Y., Duanmu, D., 2020. Natural variation at OsCERK1 regulates arbuscular mycorrhizal symbiosis in rice. New Phytologist 225, 1762–1776. https://doi.org/10.1111/nph.16158

33. Jaouad, M., Moinina, A., Ezrari, S., Lahlali, R., 2020. Key pests and diseases of citrus trees with emphasis on root rot diseases: An overview. Moroccan Journal of Agricultural Sciences 1.

34. Jeworutzki, E., Roelfsema, M.R.G., Anschütz, U., Krol, E., Elzenga, J.T.M., Felix, G., Boller, T., Hedrich, R., Becker, D., 2010. Early signaling through the Arabidopsis pattern recognition receptors FLS2 and EFR involves Ca2+-associated opening of plasma membrane anion channels. The Plant Journal 62, 367–378. https://doi.org/10.1111/j.1365-313X.2010.04155.x

35. Kaku, H., Nishizawa, Y., Ishii-Minami, N., Akimoto-Tomiyama, C., Dohmae, N., Takio, K., Minami, E., Shibuya, N., 2006. Plant cells recognize chitin fragments for defense signaling through a plasma membrane receptor. PNAS 103, 11086–11091. https://doi.org/10.1073/pnas.0508882103

36. Kamatyanatti, M., Singh, S., Sekhon, B., 2021. MUTATION BREEDING IN CITRUS-A REVIEW. Plant Cell Biotechnology and Molecular Biology 22, 1–8.

37. Kubitzki, K., Kallunki, J.A., Duretto, M., Wilson, P.G., 2011. Rutaceae, in: Kubitzki, Klaus (Ed.), Flowering Plants. Eudicots: Sapindales, Cucurbitales, Myrtaceae, The Families and Genera of Vascular Plants. Springer, Berlin, Heidelberg, pp. 276–356. https://doi.org/10.1007/978-3-642-14397-7_16

38. Latado, R.R., Neto, A.T., Figueira, A., n.d. In Vivo and in Vitro Mutation Breeding of Citrus.

39. Lee, W.-S., Rudd, J.J., Hammond-Kosack, K.E., Kanyuka, K., 2014. Mycosphaerella graminicola LysM Effector-Mediated Stealth Pathogenesis Subverts Recognition Through Both CERK1 and CEBiP Homologues in Wheat. MPMI 27, 236–243. https://doi.org/10.1094/MPMI-07-13-0201-R

40. Li, W., Hartung, J.S., Levy, L., 2006. Quantitative real-time PCR for detection and identification of Candidatus Liberibacter species associated with citrus huanglongbing. Journal of Microbiological Methods 66, 104–115. https://doi.org/10.1016/j.mimet.2005.10.018

41. Liao, D., Sun, X., Wang, N., Song, F., Liang, Y., 2018. Tomato LysM Receptor-Like Kinase SlLYK12 Is Involved in Arbuscular Mycorrhizal Symbiosis. Frontiers in Plant Science 9.

42. Liu, Y., Heying, E., Tanumihardjo, S.A., 2012. History, Global Distribution, and Nutritional Importance of Citrus Fruits. Comprehensive Reviews in Food Science and Food Safety 11, 530–545. https://doi.org/10.1111/j.1541-4337.2012.00201.x

43. Livak, K.J., Schmittgen, T.D., 2001. Analysis of relative gene expression data using real-time quantitative PCR and the 2(-Delta Delta C(T)) Method. Methods 25, 402–408. https://doi.org/10.1006/meth.2001.1262

44. Ma, W., Pang, Z., Huang, X., Xu, J., Pandey, S.S., Li, J., Achor, D.S., Vasconcelos, F.N.C., Hendrich, C., Huang, Y., Wang, W., Lee, D., Stanton, D., Wang, N., 2022. Citrus Huanglongbing is a pathogen-triggered immune disease that can be mitigated with antioxidants and gibberellin. Nat Commun 13, 529. https://doi.org/10.1038/s41467-022-28189-9

45. Magalhães, D.M., Scholte, L.L.S., Silva, N.V., Oliveira, G.C., Zipfel, C., Takita, M.A., De Souza, A.A., 2016. LRR-RLK family from two Citrus species: genome-wide identification and evolutionary aspects. BMC Genomics 17, 623. https://doi.org/10.1186/s12864-016-2930-9

46. Meng, X., Zhang, S., 2013. MAPK Cascades in Plant Disease Resistance Signaling. Annual Review of Phytopathology 51, 245–266. https://doi.org/10.1146/annurev-phyto-082712-102314

47. Miya, A., Albert, P., Shinya, T., Desaki, Y., Ichimura, K., Shirasu, K., Narusaka, Y., Kawakami, N., Kaku, H., Shibuya, N., 2007. CERK1, a LysM receptor kinase, is essential for chitin elicitor signaling in Arabidopsis. PNAS 104, 19613–19618. https://doi.org/10.1073/pnas.0705147104

48. Miyata, K., Kozaki, T., Kouzai, Y., Ozawa, K., Ishii, K., Asamizu, E., Okabe, Y., Umehara, Y., Miyamoto, A., Kobae, Y., Akiyama, K., Kaku, H., Nishizawa, Y., Shibuya, N., Nakagawa, T., 2014. The Bifunctional Plant Receptor, OsCERK1, Regulates Both Chitin-Triggered Immunity and Arbuscular Mycorrhizal Symbiosis in Rice. Plant and Cell Physiology 55, 1864–1872. https://doi.org/10.1093/pcp/pcu129

49. Morton, C.M., 2009. Phylogenetic relationships of the Aurantioideae (Rutaceae) based on the nuclear ribosomal DNA ITS region and three noncoding chloroplast DNA regions, atpB-rbcL spacer, rps16, and trnL-trnF. Organisms Diversity & Evolution 9, 52–68. https://doi.org/10.1016/j.ode.2008.11.001

50. Nagano, Y., Mimura, T., Kotoda, N., Matsumoto, R., Nagano, A.J., Honjo, M.N., Kudoh, H., Yamamoto, M., 2018. Phylogenetic relationships of Aurantioideae (Rutaceae) based on RAD-Seq. Tree Genetics & Genomes 14, 6. https://doi.org/10.1007/s11295-017-1223-z

51. Newman, M.-A., Sundelin, T., Nielsen, J., Erbs, G., 2013. MAMP (microbe-associated molecular pattern) triggered immunity in plants. Frontiers in Plant Science 4.

52. Ngou, B.P.M., Ding, P., Jones, J.D.G., 2022. Thirty years of resistance: Zig-zag through the plant immune system. The Plant Cell koac041. https://doi.org/10.1093/plcell/koac041

53. Pang, Z., Zhang, L., Coaker, G., Ma, W., He, S.-Y., Wang, N., 2020. Citrus CsACD2 Is a Target of Candidatus Liberibacter Asiaticus in Huanglongbing Disease. Plant Physiology 184, 792–805. https://doi.org/10.1104/pp.20.00348

54. Peng, A., Zou, X., He, Y., Chen, S., Liu, X., Zhang, J., Zhang, Q., Xie, Z., Long, J., Zhao, X., 2021. Overexpressing a NPR1-like gene from Citrus paradisi enhanced Huanglongbing resistance in C. sinensis. Plant Cell Rep 40, 529–541. https://doi.org/10.1007/s00299-020-02648-3

55. Pfund, C., Tans-Kersten, J., Dunning, F.M., Alonso, J.M., Ecker, J.R., Allen, C., Bent, A.F., 2004. Flagellin Is Not a Major Defense Elicitor in Ralstonia solanacearum Cells or Extracts Applied to Arabidopsis thaliana. MPMI 17, 696–706. https://doi.org/10.1094/MPMI.2004.17.6.696

56. Plieth, C., 2018. Redox Modulators Determine Luminol Luminescence Generated by Porphyrin-Coordinated Iron and May Repress “Suicide Inactivation.” ACS Omega 3, 12295–12303. https://doi.org/10.1021/acsomega.8b01261

57. Ramadugu, C., Keremane, M.L., Halbert, S.E., Duan, Y.P., Roose, M.L., Stover, E., Lee, R.F., 2016. Long-Term Field Evaluation Reveals Huanglongbing Resistance in Citrus Relatives. Plant Disease 100, 1858–1869. https://doi.org/10.1094/PDIS-03-16-0271-RE

58. Reece, P.C., Swingle, W.T., 1967. The botany of Citrus and its wild relatives. The citrus industry 1, 190–430.

59. Robatzek, S., Bittel, P., Chinchilla, D., Köchner, P., Felix, G., Shiu, S.-H., Boller, T., 2007. Molecular identification and characterization of the tomato flagellin receptor LeFLS2, an orthologue of Arabidopsis FLS2 exhibiting characteristically different perception specificities. Plant Mol Biol 64, 539–547. https://doi.org/10.1007/s11103-007-9173-8

60. Roberts, R., Mainiero, S., Powell, A.F., Liu, A.E., Shi, K., Hind, S.R., Strickler, S.R., Collmer, A., Martin, G.B., 2019. Natural variation for unusual host responses and flagellin-mediated immunity against Pseudomonas syringae in genetically diverse tomato accessions. New Phytologist 223, 447–461. https://doi.org/10.1111/nph.15788

61. Robertson, C.J., Zhang, X., Gowda, S., Orbović, V., Dawson, W.O., Mou, Z., 2018. Overexpression of the Arabidopsis NPR1 protein in citrus confers tolerance to Huanglongbing. Journal of Citrus Pathology 5. https://doi.org/10.5070/C451038911

62. Ruan, J., Li, H., 2020. Fast and accurate long-read assembly with wtdbg2. Nat Methods 17, 155–158. https://doi.org/10.1038/s41592-019-0669-3

63. Saijo, Y., Loo, E.P., Yasuda, S., 2018. Pattern recognition receptors and signaling in plant– microbe interactions. The Plant Journal 93, 592–613. https://doi.org/10.1111/tpj.13808

64. Shi, Q., Febres, V.J., Jones, J.B., Moore, G.A., 2016. A survey of FLS2 genes from multiple citrus species identifies candidates for enhancing disease resistance to Xanthomonas citri ssp. citri. Horticulture Research 3, 16022. https://doi.org/10.1038/hortres.2016.22

65. Shi, Q., Febres, V.J., Jones, J.B., Moore, G.A., 2015. Responsiveness of different citrus genotypes to the Xanthomonas citri ssp. citri-derived pathogen-associated molecular pattern (PAMP) flg22 correlates with resistance to citrus canker. Molecular Plant Pathology 16, 507–520. https://doi.org/10.1111/mpp.12206

66. Shi, Q., Febres, V.J., Zhang, S., Yu, F., McCollum, G., Hall, D.G., Moore, G.A., Stover, E., 2018. Identification of Gene Candidates Associated with Huanglongbing Tolerance, Using ‘Candidatus Liberibacter asiaticus’ Flagellin 22 as a Proxy to Challenge Citrus. MPMI 31, 200–211. https://doi.org/10.1094/MPMI-04-17-0084-R

67. Shimizu, T., Nakano, T., Takamizawa, D., Desaki, Y., Ishii-Minami, N., Nishizawa, Y., Minami, E., Okada, K., Yamane, H., Kaku, H., Shibuya, N., 2010. Two LysM receptor molecules, CEBiP and OsCERK1, cooperatively regulate chitin elicitor signaling in rice. Plant J 64, 204–214. https://doi.org/10.1111/j.1365-313X.2010.04324.x

68. Stover, E., McCollum, G., Ramos, J., Jr, R.G.S., 2014. Growth, Health and Liberibacter asiaticus Titer in Diverse Citrus Scions on Mandarin versus Trifoliate Hybrid Rootstocks in a Field Planting with Severe Huanglongbing. Proceedings of the Florida State Horticultural Society 127, 53–59.

69. Takai, R., Isogai, A., Takayama, S., Che, F.-S., 2008. Analysis of Flagellin Perception Mediated by flg22 Receptor OsFLS2 in Rice. MPMI 21, 1635–1642. https://doi.org/10.1094/MPMI-21-12-1635

70. Trdá, L., Fernandez, O., Boutrot, F., Héloir, M.-C., Kelloniemi, J., Daire, X., Adrian, M., Clément, C., Zipfel, C., Dorey, S., Poinssot, B., 2014. The grapevine flagellin receptor VvFLS2 differentially recognizes flagellin-derived epitopes from the endophytic growth-promoting bacterium Burkholderia phytofirmans and plant pathogenic bacteria. New Phytologist 201, 1371–1384. https://doi.org/10.1111/nph.12592

71. USDA, National Agricultural Statistics Service. 2020. Citrus Fruits: 2020 Summary.

72. Uzun, A., Yesiloglu, T., 2012. Genetic Diversity in Citrus. InTech.

73. Veluchamy, S., Hind, S.R., Dunham, D.M., Martin, G.B., Panthee, D.R., 2014. Natural Variation for Responsiveness to flg22, flgII-28, and csp22 and Pseudomonas syringae pv. tomato in Heirloom Tomatoes. PLOS ONE 9, e106119. https://doi.org/10.1371/journal.pone.0106119

74. Vetter, M.M., Kronholm, I., He, F., Häweker, H., Reymond, M., Bergelson, J., Robatzek, S., de Meaux, J., 2012. Flagellin Perception Varies Quantitatively in Arabidopsis thaliana and Its Relatives. Molecular Biology and Evolution 29, 1655–1667. https://doi.org/10.1093/molbev/mss011

75. Wang, L., Albert, M., Einig, E., Fürst, U., Krust, D., Felix, G., 2016. The pattern-recognition receptor CORE of Solanaceae detects bacterial cold-shock protein. Nature Plants 2, 1–9. https://doi.org/10.1038/nplants.2016.185

76. Wang, N., 2019. The Citrus Huanglongbing Crisis and Potential Solutions. Molecular Plant 12, 607–609. https://doi.org/10.1016/j.molp.2019.03.008

77. Wang, P., Zhou, L., Jamieson, P., Zhang, L., Zhao, Z., Babilonia, K., Shao, W., Wu, L., Mustafa, R., Amin, I., Diomaiuti, A., Pontiggia, D., Ferrari, S., Hou, Y., He, P., Shan, L., 2020. The Cotton Wall-Associated Kinase GhWAK7A Mediates Responses to Fungal Wilt Pathogens by Complexing with the Chitin Sensory Receptors. The Plant Cell 32, 3978–4001. https://doi.org/10.1105/tpc.19.00950

78. Wang, X., Xu, Y., Zhang, S., Cao, L., Huang, Y., Cheng, J., Wu, G., Tian, S., Chen, C., Liu, Y., Yu, H., Yang, X., Lan, H., Wang, N., Wang, L., Xu, J., Jiang, X., Xie, Z., Tan, M., Larkin, R.M., Chen, L.-L., Ma, B.-G., Ruan, Y., Deng, X., Xu, Q., 2017. Genomic analyses of primitive, wild and cultivated citrus provide insights into asexual reproduction. Nat Genet 49, 765–772. https://doi.org/10.1038/ng.3839

79. Wei, Y., Balaceanu, A., Rufian, J.S., Segonzac, C., Zhao, A., Morcillo, R.J.L., Macho, A.P., 2020. An immune receptor complex evolved in soybean to perceive a polymorphic bacterial flagellin. Nat Commun 11, 3763. https://doi.org/10.1038/s41467-020-17573-y

80. Wu, G.A., Terol, J., Ibanez, V., López-García, A., Pérez-Román, E., Borredá, C., Domingo, C., Tadeo, F.R., Carbonell-Caballero, J., Alonso, R., Curk, F., Du, D., Ollitrault, P., Roose, M.L., Dopazo, J., Gmitter, F.G., Rokhsar, D.S., Talon, M., 2018. Genomics of the origin and evolution of Citrus. Nature 554, 311–316. https://doi.org/10.1038/nature25447

81. Xue, D.-X., Li, C.-L., Xie, Z.-P., Staehelin, C., 2019. LYK4 is a component of a tripartite chitin receptor complex in Arabidopsis thaliana. Journal of Experimental Botany 70, 5507–5516. https://doi.org/10.1093/jxb/erz313

82. Yu, Y., Chen, C., Huang, M., Yu, Q., Du, D., Mattia, M.R., Gmitter, F.G., 2018. Genetic Diversity and Population Structure Analysis of Citrus Germplasm with Single Nucleotide Polymorphism Markers. Journal of the American Society for Horticultural Science 143, 399–408. https://doi.org/10.21273/JASHS04394-18

83. Zhang, W., Fraiture, M., Kolb, D., Löffelhardt, B., Desaki, Y., Boutrot, F.F.G., Tör, M., Zipfel, C., Gust, A.A., Brunner, F., 2013. Arabidopsis RECEPTOR-LIKE PROTEIN30 and Receptor-Like Kinase SUPPRESSOR OF BIR1-1/EVERSHED Mediate Innate Immunity to Necrotrophic Fungi. The Plant Cell 25, 4227–4241. https://doi.org/10.1105/tpc.113.117010

84. Zhang, X., Henriques, R., Lin, S.-S., Niu, Q.-W., Chua, N.-H., 2006. Agrobacterium-mediated transformation of Arabidopsis thaliana using the floral dip method. Nat Protoc 1, 641–646. https://doi.org/10.1038/nprot.2006.97

85. Zhong, G., Nicolosi, E., 2020. Citrus Origin, Diffusion, and Economic Importance, in: Gentile, A., La Malfa, S., Deng, Z. (Eds.), The Citrus Genome, Compendium of Plant Genomes. Springer International Publishing, Cham, pp. 5–21. https://doi.org/10.1007/978-3-030-15308-3_2

